# Self-Supervised AI Reveals a Hidden Landscape of Prognostic Spatial Patterns in Multiplex Immunofluorescence Images

**DOI:** 10.1101/2025.10.16.682563

**Authors:** Lucas Farndale, Kai Rakovic, Ian R. Powley, Phimmada Hatthakarnkul, Farzaneh Seyedshahi, Sona Szalma, Leonor Schubert Santana, Silvia Martinelli, Fiona Ballantyne, Rachel L. Baird, Leah Officer-Jones, David Moore, Claire R. Wilson, Marco Sereno, Noori Maka, Campbell Roxburgh, Crispin J. Miller, David Chang, Joanne Edwards, Edward W. Roberts, John Le Quesne, Ke Yuan

**Affiliations:** School of Cancer Sciences, University of Glasgow, Glasgow, United Kingdom; Cancer Research UK Scotland Institute, Glasgow, United Kingdom; Department of Pathology, Queen Elizabeth University Hospital, Glasgow, United Kingdom; School of Medicine, Medical Sciences and Nutrition, University of Aberdeen, Aberdeen, United Kingdom; Research Department of Oncology, University College London, London, United Kingdom; Leicester Medical School, University of Leicester, Leicester, United Kingdom; Division of Respiratory Sciences, School of Medical Sciences, University of Leicester, Leicester, United Kingdom; Academic Unit of Surgery, University of Glasgow, Glasgow, United Kingdom

## Abstract

Modern spatial proteomic methods, such as multiplex immunofluorescence (mIF) imaging, offer a data-rich view of spatial biology in intact tissues. However, interpreting its complexity is a major bottleneck, limiting its potential for biological discovery and clinical translation. Current computational methods often rely solely on segmentation-based analysis that discards crucial morphological information and are limited to testing pre-defined hypotheses. Here, we introduce a self-supervised learning (SSL) framework, HyPERSTAC, which integrates SSL and segmentation-based analysis to enable hypothesis-agnostic, context-aware discovery of biomarkers directly from mIF images. Our approach extracts rich feature representations that capture holistic architectural patterns, which integrate cellular morphology, marker interactions, and microenvironmental context without human supervision. Applying HyPERSTAC to close to 400,000 mIF tissue images from two cancer cohorts totalling over 1,800 patients, we show that SSL-derived features complement existing spatial analyses. Using HyPERSTAC to integrate SSL-derived features with segmentation-based neighbourhood analysis or the established spatial frameworks *TRACERx-PHLEX* and *MuSpAn* increased prognostic concordance in held-out patients. HyPERSTAC improves survival prediction for four SSL training methods, and two state-of-the-art foundation models. The method autonomously identified previously unknown and potentially clinically actionable biological patterns. In lung adenocarcinoma, these include a Ki67-mediated immune evasion phenotype, a subcellular pattern of GLB1 expression which aligns with low-grade *EGFR*-driven tumours, and distinct modes of tumour-immune interaction in PD-L1^+^ patients.

We also find a regulatory T-cell-mediated immuno-suppressive environment promoting tumour budding in colorectal carcinoma. Our work shows that SSL can complement segmentation-based analysis, establishing HyPERSTAC as a powerful, scalable, explainable, and unbiased platform to decode tissue ecosystems without pre-defined hypotheses. This transforms high-plex imaging from a hypothesis-testing tool into a hypothesis-generating engine that can accelerate the discovery of next-generation spatial biomarkers. We release two large image datasets from *LATTICe-A*, creating the largest publicly available multiplex immunofluorescence patient cohort, with two panels totalling almost 6,000 images from 1,000 patients and accompanying survival data. This creates a new benchmark for the community and enables further biomarker discovery from these data.

## 1 Main

Multiplex immunofluorescence (mIF) allows simultaneous quantification of many protein markers within a single intact tissue section, enabling spatially informed precision oncology [1]. However, the richness of mIF data has outpaced our ability to interpret it, leaving its transformative potential unrealised. Clinical histopathology is routinely supplemented by additional single-plex immunohistochemical protein detection assays to help guide patient management, for example to select patients for personalised therapies in lung adenocarcinoma (LUAD) [2, 3, 4] or to augment diagnosis in colorectal carcinoma (CRC) [5]. Sequential single-plex assays are tissue-exhaustive, a key limitation when personalised therapies and associated biomarkers proliferate [6] as diagnostic tissue is a scarce resource.

Widespread use of mIF faces two fundamental analytical challenges. First, human perception is ill-equipped to interpret high-plex images: simultaneously viewing dozens of markers exceeds cognitive limits, and composite visualisations become crowded, obscuring critical interactions. Second, conventional computational tools rely on cell segmentation, reducing complex tissue ecosystems to simplistic cell-type counts or neighbourhood graphs [7, 8]. While offering valuable insights for explicit hypothesis testing, segmentation pipelines discard morphological information and, crucially, discovery is typically confined to pre-defined, manually generated hypotheses. Even with perfect segmentation, the combinatorial explosion of cell states and interactions makes exhaustive analysis prohibitive due to the sample size required for significance with this number of comparisons.

Established segmentation-based spatial frameworks have made such analyses reproducible and scalable. *TRACERx-PHLEX* combines deep-learning cell segmentation and automated phenotyping with interpretable spatial metrics such as cellular barrier scores [9], and *MuSpAn* provides spatial statistics, topological data analysis and network methods within a unified toolbox [10]. Both quantify relationships between predefined cell types, so what they can detect is bounded by segmentation and phenotyping and by the relationships the analyst chooses to measure. Segmentation-free approaches remove this dependence. In spatial transcriptomics, *FICTURE* infers pixel-level spatial factors directly from submicron-resolution transcript coordinates [11]. In multiplexed imaging, *Pixie* clusters pixels to annotate features that lie outside cells, such as the extracellular matrix [12], and *CANVAS* uses self-supervised masked image modelling to learn representations of tumour microenvironments without segmentation [13].

Self-supervised learning (SSL) offers a paradigm shift by learning directly from image data without manual supervision. In contrast to supervised methods constrained by annotated training sets [14, 15] joint-embedding SSL extracts semantically meaningful features by identifying invariant patterns across augmented views of the same image. Typically, redundant patterns such as biological noise are obscured by the augmentations, while meaningful patterns such as protein localisation are unaffected [16, 17]. This approach is uniquely well-suited to mIF: it preserves spatial and morphological information, integrates signals across all markers simultaneously, and avoids errors and biases introduced by segmentation or pre-defined cell states. SSL applications in pathology have focused on brightfield images [18, 19, 20, 21, 22], but the potential to discover high-order, cross-protein spatial biomarkers in full mIF datasets remains untapped. Our framework, HyPERSTAC (**Hy**pothesis-free **P**roteomic **E**mbeddings for **R**epresenting **S**patial **T**issue **A**rchitecture & **C**ontext), incorporates both segmentation-based analysis and SSL, enabling deterministic, quantified cell-based analysis while learning rich spatial features directly from image pixels. We then use learned image features as a substrate for Leiden community detection, discovering spatial proteomic clusters (SPCs) - clusters of similar image tiles that represent recurrent spatial neighbourhoods. Through HyPERSTAC’s unique interplay between segmentation and SSL, mIF images are characterised using traditional segmentation for interpretation, and SSL for discovery, bridging the gap between data-driven discovery and established pathology.

The key contribution of this work is that we are able to use HyPERSTAC to discover spatial biomarkers by holistically considering tissue architecture and cell interactions while retaining information about their morphology or marker distribution. No existing study has enabled hypothesis-agnostic discovery of candidate biomarkers from mIF which have withstood validation in holdout patient sets. While recent work has applied SSL to multiplexed imaging [14, 23, 13, 24, 25], these approaches are typically developed on small cohorts, are generally used in place of rather than alongside segmentation-based analysis, and focus on characterising cells or patients rather than holistic tissue morphology. They have rarely been evaluated on held-out patients from large cohorts or benchmarked against established spatial frameworks. HyPERSTAC is distinctive in integrating SSL with segmentation-based analysis, and in evaluating this integration in held-out patients from large cohorts against established spatial frameworks. Here, we introduce a discriminative SSL framework that identifies prognostic biomarkers from 398,502 tissue microarray (TMA) images from over 6,000 individual cores across two cancers. We release these images and accompanying survival data to the community, contributing the largest public dataset of multiplex immunofluorescence images, as well as a benchmark for survival prediction using spatial proteomics foundation models on a large patient cohort. HyPERSTAC identifies prognostic tissue patterns, integrating marker interactions, cellular morphology, and tumour microenvironment context without prior assumptions. We evaluate the method in four ways: (i) predicting survival in held-out patients, benchmarked against segmentation-based neighbourhood analysis, the established spatial frameworks TRACERx-PHLEX and MuSpAn (Fig. 2b; Supplementary Fig. S5), and six SSL encoders including two multiplex foundation models; (ii) testing whether HyPERSTAC risk scores are prognostic independently of clinicopathological factors; (iii) uncovering novel candidate biomarkers, and (iv) showing cross-panel generality through integrated analysis of independent staining panels. Crucially, all biomarkers emerge from the data itself — without manual supervision or pre-formed hypotheses — yet remain interpretable through pathologist-validated patterns.

## 2 Results

A typical, hypothesis-driven mIF analysis would involve cell segmentation followed by assignment of phenotypes to cells using the fluorescence intensities of key lineage markers. We propose HyPERSTAC, allowing the SSL model to identify key recurrent image features which are integrated with the classical segmentation-based approach. This allows us to leverage both the transparent, well-quantified features of classical analysis, and the holistic, qualitative features of the raw image, to maximise the model’s information usage. We used three mIF panels from two common solid malignancies to demonstrate the utility of our methodology. The Leicester Archival Thoracic Tumour Investigatory Cohort - Adenocarcinoma (LATTICe-A) cohort [26] was stained with two panels: an immuno-oncology (LATTICe-A-IO) panel, and a hallmarks of cancer panel (LATTICe-A-Hallmarks). We also used an immuno-oncology panel on cores from the Glasgow Royal Infirmary (GRI) colorectal cancer cohort [27]. The details of the constituents of each panel are detailed in Supplementary Figure S1 and in the methods.

Initially, we aggregate cell lineage densities per case, and use it to predict prognosis. In the LATTICe-A-IO panel, this gives us a modest ability to stratify patients by CD8^+^ T-cell density (Supplementary Figure S2a, b). We also perform classical neighbourhood discovery on all panels, which achieves modest prognostic performance (Figure 2b); below, we benchmark HyPERSTAC against this and two other established spatial analysis frameworks, *TRACERx-PHLEX* and *MuSpAn* (Supplementary Figure S5).

HyPERSTAC generates patterns of interest from 224 *×* 224px tiles from each core based on their latent representations, and incorporates segmentation-based cell phenotypes to enable their interpretation. This means that for each tile we were able to quantify the SSL-discovered phenotypes by classically discovered cell types, aiding the interpretation of the SPCs. These SPC frequencies were then linked to outcome and other key clinicopathological states (Figure 1).

**Figure 1.**
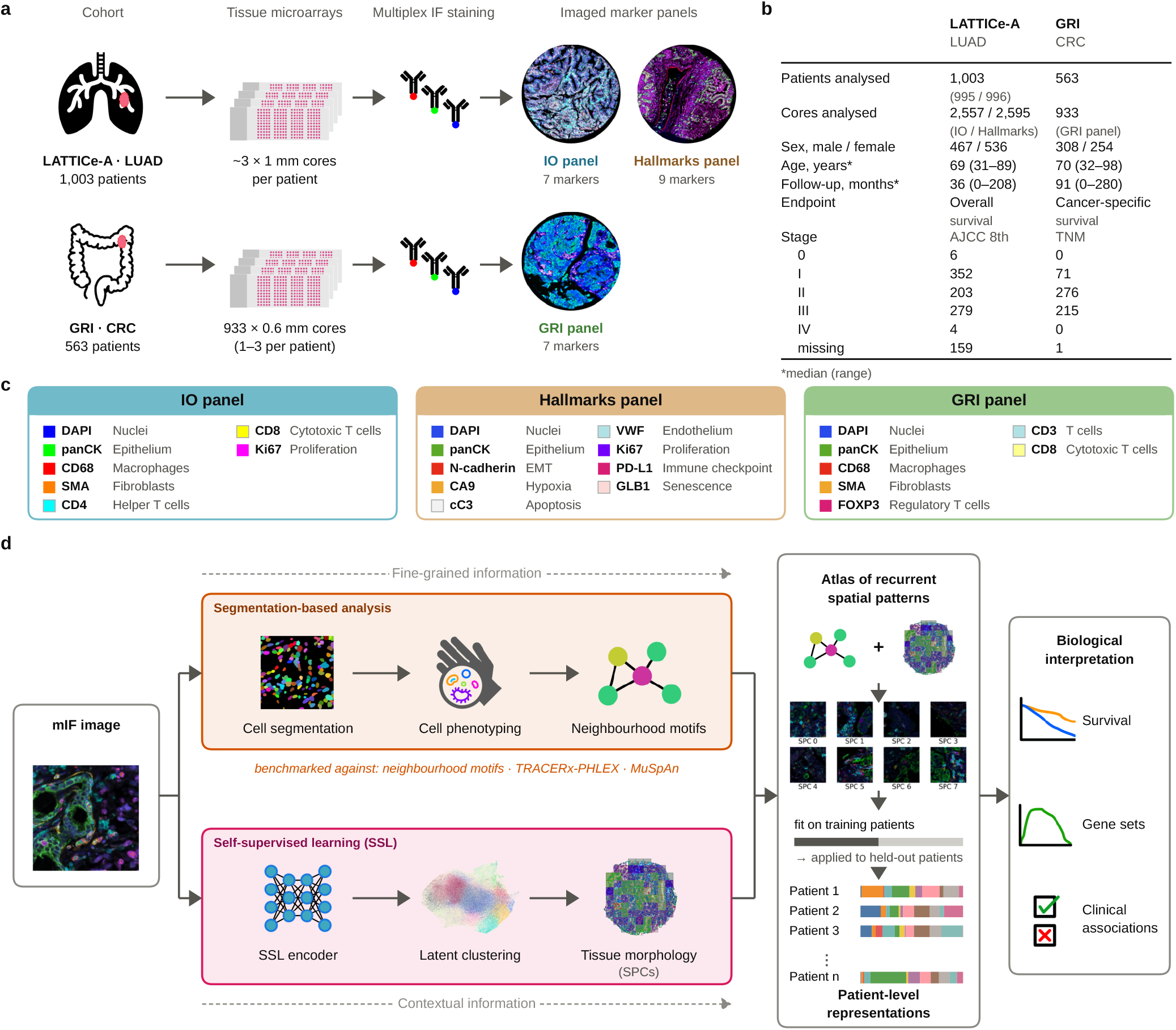
HyPERSTAC integrates self-supervised and segmentation-based analysis to decode spatial proteomic patterns from multiplex immunofluorescence cohorts. (a) Cohorts and imaging: LATTICe-A lung adenocarcinoma (LUAD; 1,003 patients) stained with the immuno-oncology (IO) and hallmarks of cancer (Hallmarks) multiplex immunofluorescence panels, and GRI colorectal carcinoma (CRC; 563 patients) stained with an immune panel. (b) Characteristics of the analysed patients (median (range) for age and follow-up; GRI sex and age are unknown for one patient each). (c) Markers in each panel and the pseudocolour used in all images. (d) Workflow. The segmentation route (orange) phenotypes cells and clusters their neighbourhoods into motifs, the information used by established frameworks such as TRACERx-PHLEX and MuSpAn. The self-supervised route (magenta) clusters tile embeddings (111 *µ*m tiles) into spatial proteomic clusters (SPCs), which capture tissue context that is not reduced to cell counts. Clusters are fitted on training patients and applied to held-out patients, and each patient is described by the frequency of each motif and SPC.

### 2.1 Automated discovery of recurrent prognostic spatial clusters

HyPERSTAC identifies recurrent spatial patterns across the data, encoding these features into SPCs. For each mIF panel, we annotate each SPC with a short description based on a visual histopathological assessment from expert pathologists (Figure 3a, c). We saw broad co-localisation of major cell lineages in distinct locales on the uniform manifold approximation and projection (UMAP) plot (Figure 3a, Supplementary Figures S3a). We saw similar patterns replicated across the LATTICe-A-Hallmarks panel (Figure 3b, Supplementary Figures S3b).

**Figure 2.**
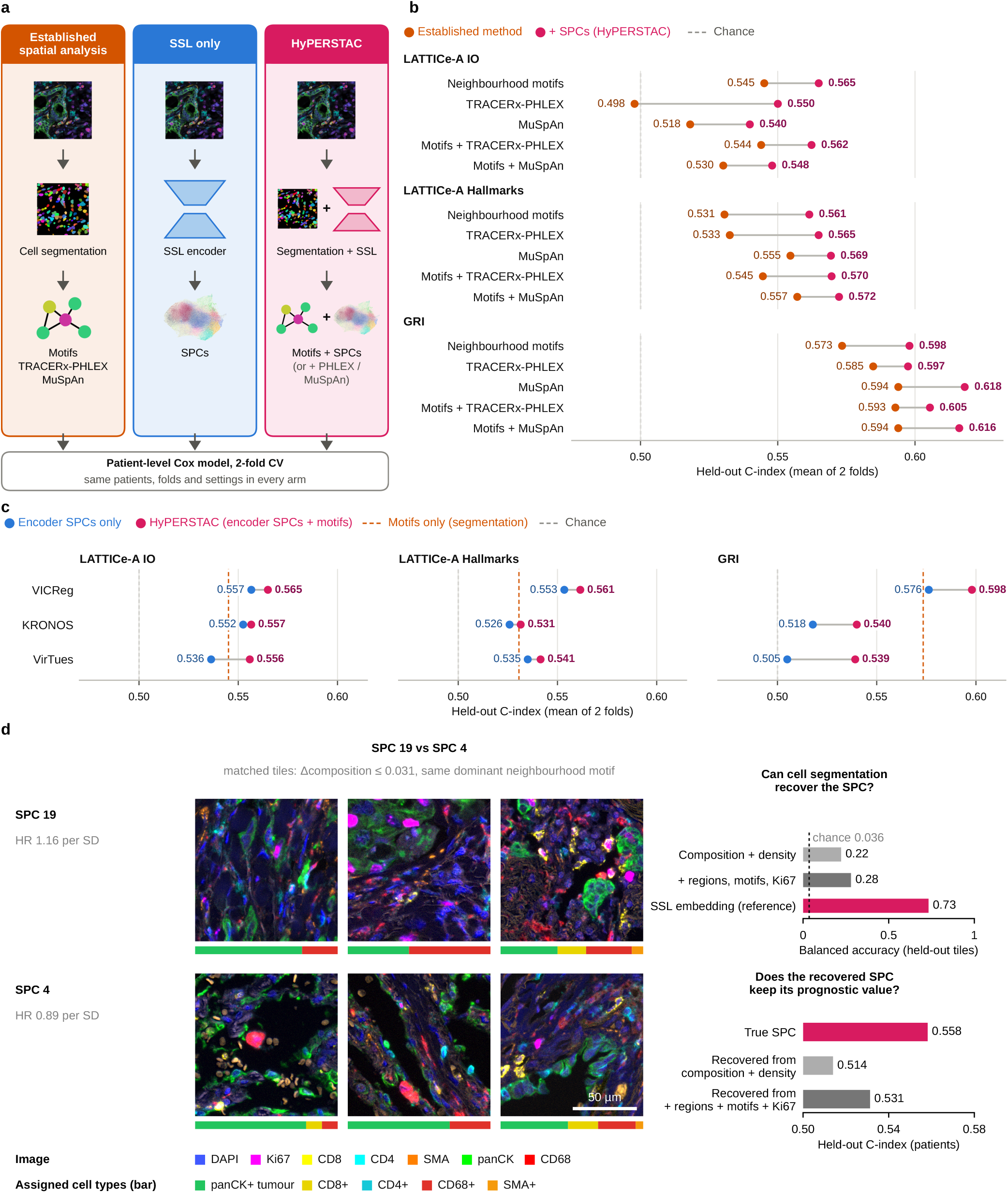
HyPERSTAC outperforms SSL and segmentation baselines across multiple encoders and cohorts. (a) Feature sets compared, each evaluated with the same patient-level Cox model, patients and two-fold cross-validation. (b) Held-out concordance index (C-index) of each established method alone (orange) and with SPCs added (HyPERSTAC; magenta) in LATTICe-A-IO (*n* = 995), LATTICe-A-Hallmarks (*n* = 996) and GRI (*n* = 563). A greater C-index means the SPCs added information; adding SPCs raised C for every method in every cohort. Dashed line, chance (C=0.5). (c) Robustness to the encoder. Held-out C of SPCs alone (blue) and HyPERSTAC (magenta) for VICReg, trained on our images, and the foundation models KRONOS and VirTues (all six encoders in Supplementary Figure S6); dashed orange line, motifs alone. (d) What SPCs capture beyond cell composition. Tiles with near-identical cell-type composition (bars; assigned cell types only) and the same dominant motif, from different patients, can fall into SPCs with opposite prognostic associations (SPC 19, training-fold HR 1.16 per s.d.; SPC 4, HR 0.89): the encoder separates tissue that segmentation describes as the same. Illustrative pair selected post hoc (pre-specified pair in Supplementary Figure S7). Tiles 111 *µ*m; scale bar, 50 *µ*m. Right, segmentation features recover a held-out tile’s SPC poorly (balanced accuracy; dashed line, chance), and replacing true SPCs with recovered ones loses most of their prognostic value (*n* = 995).

**Figure 3.**
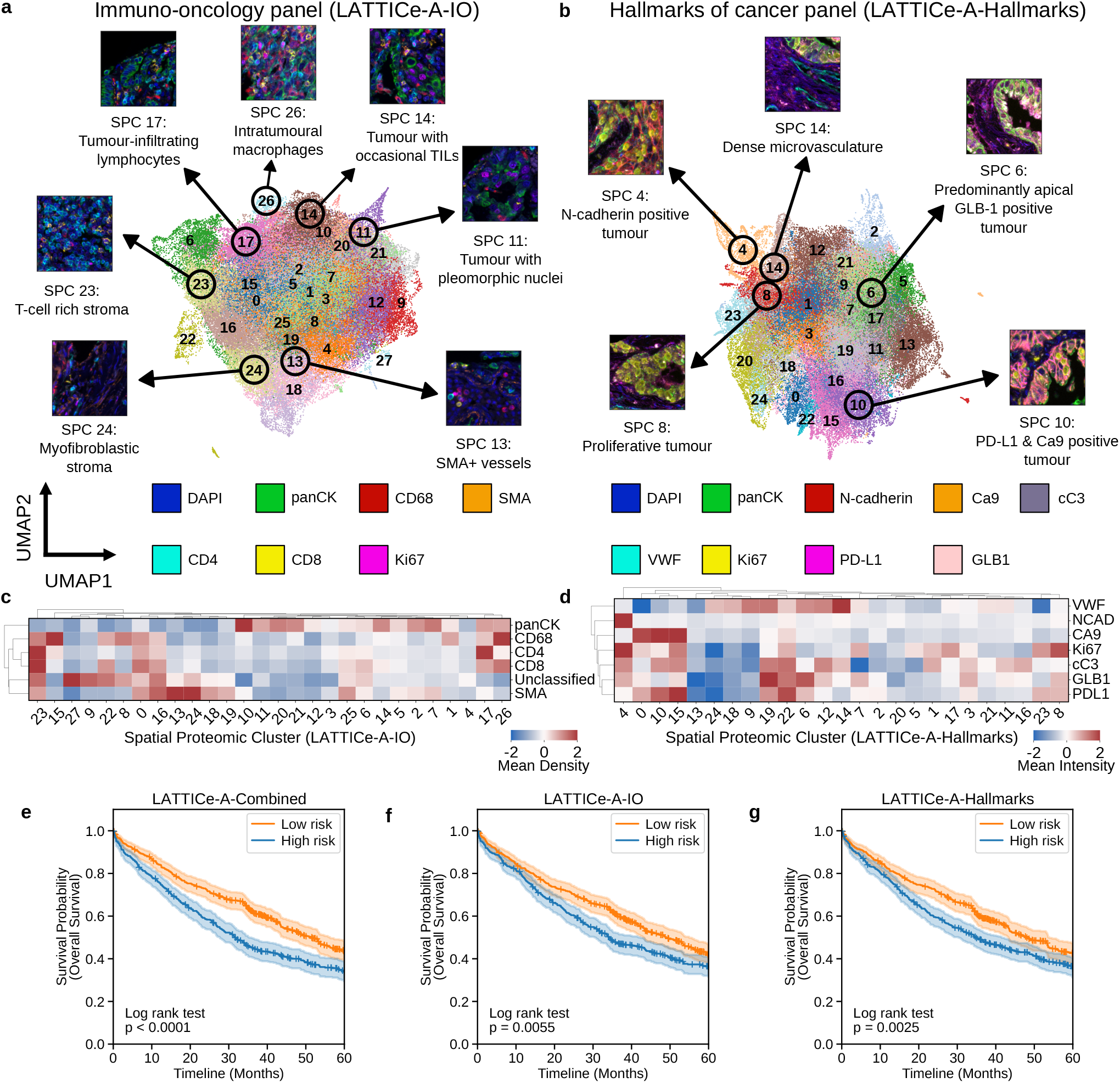
HyPERSTAC resolves tumour, immune, and stromal microenvironments into reproducible spatial proteomic clusters with distinct cell compositions. (a, b) UMAP of the LATTICe-A-IO (a) and LATTICe-A-Hallmarks (b) tile embeddings, coloured by SPC, with example tiles (111 *µ*m) and pathologist descriptions. Tumour, stroma and immune tissue occupy separate regions, and SPCs resolve recurrent arrangements within them. Supplementary Figure S3 colours the same embeddings by each marker. (c, d) Mean density of each cell type (c, IO) and mean intensity of each marker (d, Hallmarks) per SPC, as z-scores: most SPCs have a distinct composition. (e–g) Overall survival of held-out patients split into high (blue) and low (orange) risk by SPC frequencies alone (VICReg; no segmentation features), using both panels (e; *n* = 1,003; log-rank *p* = 1.8 *×* 10^*−*5^), IO (f; *n* = 995; *p* = 0.0055) or Hallmarks (g; *n* = 996; *p* = 0.0025). Risk cut-points were set on training patients only. Shading, 95% CI; +, censored.

Despite the clear segregation of immune-, stromal- and tumour-dominated regions in the latent space, in the LATTICe-A-IO panel we identified that heterogeneous higher-order interactions between cells are captured by HyPERSTAC’s latent representations. For example, we saw clear division in the latent space between SPCs with and without inflammatory cells, as well as neighbourhoods characterised by recurrent relative compositions of these cells (Figure 3a, c, Supplementary FigureS3a). Similarly, in the LATTICe-A-Hallmarks panel we saw SPCs characterised by tumour cells expressing PD-L1, or by hypoxia (which show concomitant downregulation of angiogenesis; Figure 3d).

We used a conservative two-fold cross validation to establish the realistic prognostic power of the SPCs we discovered, yielding substantial test set sizes with which to validate our observations (larger holdout sets than standard five-fold). Our predictive model was able to successfully stratify patients into high- and low-risk groups using combined information from both panels and individually (Figure 3e-g, Supplementary Figure S8; SPC-level hazard ratios in Supplementary Figure S4).

### 2.2 Discovered clusters complement established spatial analyses

To benchmark HyPERSTAC, we evaluated every feature set with the same patient-level Cox pipeline, patients and two-fold cross-validation splits (Figure 2a; Methods). Alongside neighbourhood motifs, we used two established spatial analysis frameworks: TRACERx-PHLEX [9], which quantifies cellular barriers and spatial clustering, and MuSpAn [10], from which we pre-specified spatial statistics and neighbourhood, correlation-map and topological descriptors.

Adding SPCs increased the concordance index (C-index) of every established method in every cohort (Figure 2b, Supplementary Figure S5). The gain was significant in nine of the ten LUAD comparisons in Figure 2b (change in C-index, ΔC, +0.015 to +0.052; for example, TRACERx-PHLEX alone C-index 0.498, with SPCs 0.550, in LATTICe-A-IO) and of similar size in GRI (ΔC +0.013 to +0.025; about 62 cancer-specific deaths per test fold). In LATTICe-A-IO, HyPERSTAC (motifs + SPCs) also outperformed motifs + MuSpAn (ΔC +0.035, 95% CI +0.014 to +0.054) and motifs + TRACERx-PHLEX (ΔC +0.021, 95% CI +0.002 to +0.039), whereas neither framework added to motifs. In LATTICe-A-Hallmarks, where the motif baseline clusters neighbourhood marker intensities rather than phenotyped cells (Methods), MuSpAn also improved on motifs, and HyPERSTAC performed similarly to motifs + MuSpAn (ΔC +0.004, 95% CI *−* 0.017 to +*−* 0.027) and motifs + TRACERx-PHLEX (ΔC +0.017, 95% CI 0.006 to +0.040). Whether SPCs or segmentation-based analyses achieved better prognostic performance varied by cohort (Supplementary Figure S6), but their value lies in combination with segmentation.

To see what SPCs capture that segmentation does not, we compared LATTICe-A-IO tiles with near-identical cell-type composition and the same dominant neighbourhood motif. Such tiles could fall in SPCs with opposite prognostic associations, such as SPC 19 (compact stroma–tumour interface; training-fold HR 1.16 per s.d.) and SPC 4 (alveolar architecture; HR 0.89) (Figure 2d; Supplementary Figure S7). A classifier recovered a held-out tile’s SPC poorly from segmentation features (balanced accuracy 0.22–0.28; chance 0.036), and replacing each tile’s SPC by the one recovered from cell composition and density reduced held-out C-index from 0.558 to 0.514 (ΔC 0.044, 95% CI 0.015–0.072). Part of the signal is nevertheless carried by conventional markers: SPC 19 is enriched for Ki67^+^ cells, and adding tumour/stroma regions, motif fractions and the Ki67^+^ fraction narrowed the gap (C 0.531, p = 0.062). This illustrates that when we know the feature we are trying to characterise, it is often possible to characterise SPC-derived features using segmentation-based descriptors. However, the overwhelming number of possible combinations of descriptors means that this is an infeasible method for discovery. HyPERSTAC’s SPCs enable this discovery and hypothesis generation.

Among six encoders, four self-supervised methods trained on our images (VICReg [17], SimCLR [16], DINOv2 [28], MAE [29]) and two multiplex foundation models (KRONOS [30], VirTues [25]), VICReg ranked first (average rank 1.3; Supplementary Figure S6c). SPCs added significantly to motifs in LUAD for VICReg, SimCLR and MAE, but not for DINOv2, whose held-out LATTICe-A tiles collapsed onto a few training clusters, or for the foundation models (Supplementary Figure S6). HyPERSTAC therefore does not depend on a single encoder, but benefits most from encoders trained on the study images.

Finally, in LUAD the held-out HyPERSTAC risk score remained prognostic after adjustment for stage, IASLC grade, age and sex in LATTICe-A-Hallmarks (HR 1.19 per s.d., 95% CI 1.08–1.30; *n* = 780, 433 deaths) but marginally missed significance in LATTICe-A-IO (HR 1.09, 95% CI 0.99–1.20). In colorectal cancer, the independent effect was larger and concentrated in stage III disease (Figure 5b; see below).

### 2.3 Proliferating tumour cells evade immune attack

Several SPCs from the LATTICe-A-IO panel were characterised by high Ki67 expression in their constituent cells (Figure 4a). We sought to look at SPC 22 in closer detail, as it was characterised by proliferating tumour cells adjacent to a mixed inflammatory population (Figure 4b), but it had no clear survival association in its own right. This led us to hypothesise that T-cell density may be less beneficial for survival in the context of high tumour proliferation. Therefore we explored its clinicopathological significance, by investigating the relationship between T-cell density, proliferation and prognosis. We stratified all patients by their tumour cell mean Ki67 fluorescence intensity and T-cell density, and found that T-cell density is a positive prognostic factor only in the low-proliferation setting (Figures 4c, d). This remained in a sub-analysis of PD-L1 positive patients, which is more common in the high proliferative group (Supplementary Figures S9a-c). TP53-mutant tumours were more often highly proliferative, but the pattern was not explained by TP53 status: T-cell density was associated with longer survival in Ki67-low tumours whether TP53 was wild type or mutant, in both LATTICe-A and TCGA-LUAD, and adding TP53 to the model left the T-cell *×* Ki67 interaction unchanged (Supplementary Figure S9d, e).

**Figure 4.**
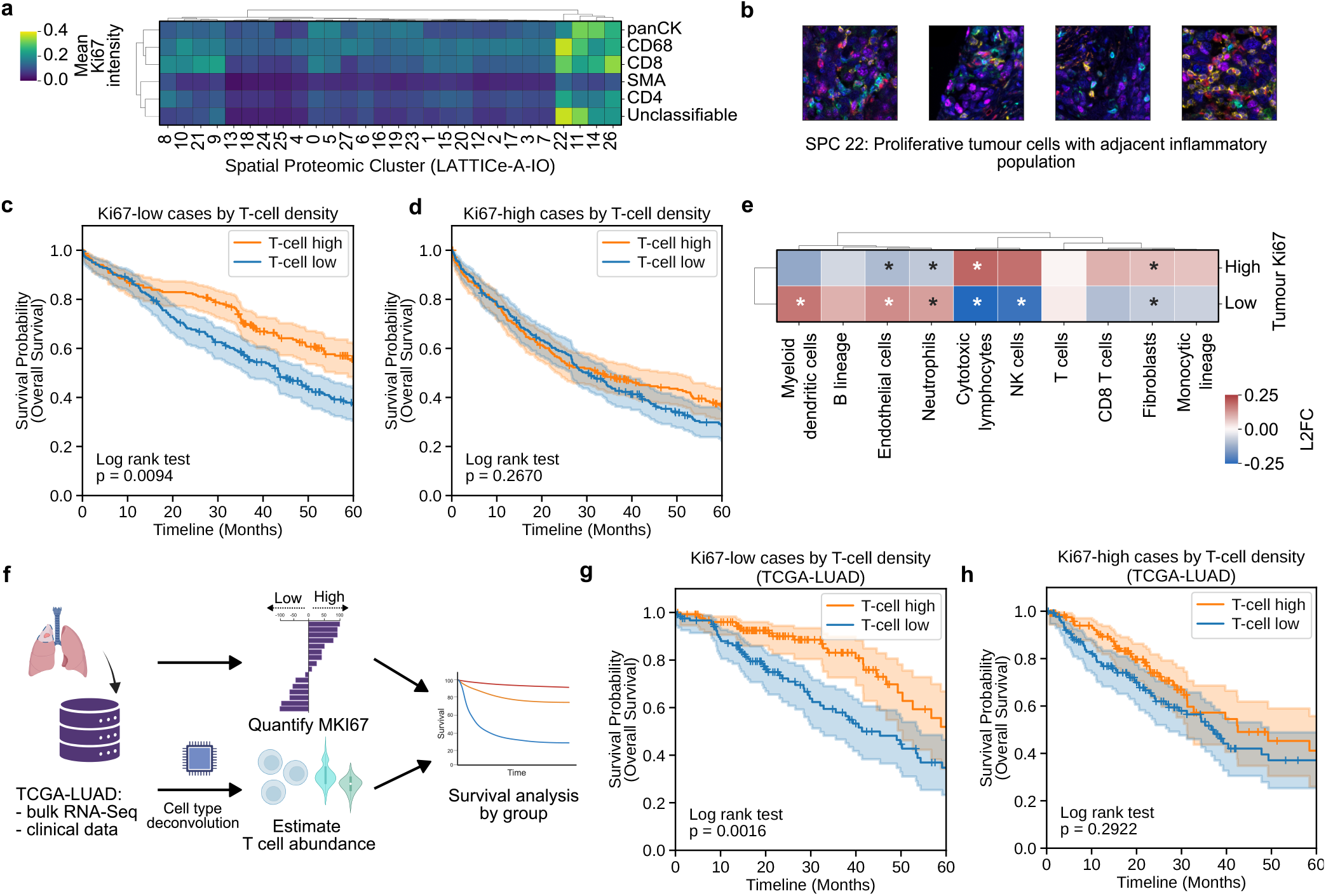
Proliferating tumour cells abrogate the prognostic benefit of T-cell infiltration in lung adenocarcinoma. (a) Mean Ki67 intensity of each cell type in each LATTICe-A-IO SPC: a few SPCs, including SPC 22, contain highly proliferating tumour cells. (b) SPC 22 tiles (111 *µ*m): proliferating tumour cells beside a mixed inflammatory infiltrate. (c, d) Overall survival by T-cell density (median split) in tumours with low (c) or high (d) tumour-cell Ki67 (median split): T cells are associated with longer survival in Ki67-low (c; *n* = 441; log-rank *p* = 0.0094) but not Ki67-high tumours (d; *n* = 481; *p* = 0.27). Associative analysis across the cohort. (e) Enrichment of MCP counter cell type deconvolution estimates in Ki67 high versus low cases expressed as log_2_ fold change. Statistical significance tested with a two-tailed Mann-Whitney-U test; * *p <* 0.05 after correction for multiple testing with the Benjamini-Hochberg method. (f)Schematic of validation on TCGA-LUAD bulk RNA-Seq data. (g, h) Replication in TCGA-LUAD, with proliferation from *MKI67* and T cells from MCP-counter: T cells are again associated with survival in *MKI67* -low (g; *n* = 252; *p* = 0.0016) but not *MKI67* -high tumours (h; *n* = 252; *p* = 0.29). Shading, 95% CI; +, censored.

Using RNA-Seq-based cell type deconvolution [31], we found small but significant differences in microenvironmental cell abundance estimates between Ki67-high/T-cell-high and Ki67-low/T-cell-high tumours. The Ki67-high tumours had increased fibroblast numbers and more cytotoxic lymphocytes compared to the Ki67-low tumours, which were better vascularised, and had increased myeloid dendritic cells and neutrophils (Figure 4e).

In order to validate this observation, we similarly classified patients using bulk transcriptomic data from The Cancer Genome Atlas (TCGA) [32]. We estimated T cell abundance using microenvironment cell type deconvolution [31] and used *MKI67* gene expression as a marker of proliferation (Figure 4f).

We found, again, that the ability of T-cell density to stratify patients into risk groups is successful only in low-proliferation tumours (Figures 4g, h). This has implications for translatability, suggesting that we should consider proliferation rates alongside measures of immune cell engagement when assessing prognosis and the likelihood of response to immunotherapies.

### 2.4 Regulatory T-cells contribute to the high-stroma phenotype in colorectal carcinoma

Next, we turned our attention to the GRI cohort. As with the LUAD panels, we found broad structure in HyPERSTAC’s tile embedding space, with predominantly-stromal and -tumour SPCs occupying separate niches (Figure 5a, Supplementary Figures S3c). We were, again, able to link the frequency of SPCs to prognosis in the held-out test patients (Supplementary Figure S8). The HyPERSTAC risk score separated patients with stage III disease into risk groups (log-rank p = 0.0048; high risk *n* = 103, 49 deaths; low risk *n* = 112, 31 deaths; Figure 5b), and across all stages it remained independently prognostic after adjustment for TNM stage, age and sex (HR 1.36 per s.d., 95% CI 1.16–1.59, p = 1.1 *×* 10^*−*4^) and after further adjustment for tumour budding, Klintrup–Mäkinen grade, tumour–stroma percentage, mismatch-repair status, and differentiation (HR 1.39, 95% CI 1.18–1.65, p = 9.2 *×* 10^*−*5^; *n* = 509, 116 deaths). Stage II patients showed no separation (log-rank p = 0.49; Supplementary Figure S11). We again calculated cell density measures for each cluster on this panel and found distinctions between stroma and tumour clusters (Figure 5c). In particular, there was marked heterogeneity in stromal composition amongst those stromal clusters. SPC 11 was notable, as this was enriched in regulatory T-cells (T_reg_) and macrophages while being depleted for CD8^+^ T-cells, suggesting a pro-tumour microenvironment (Figure 5c).

**Figure 5.**
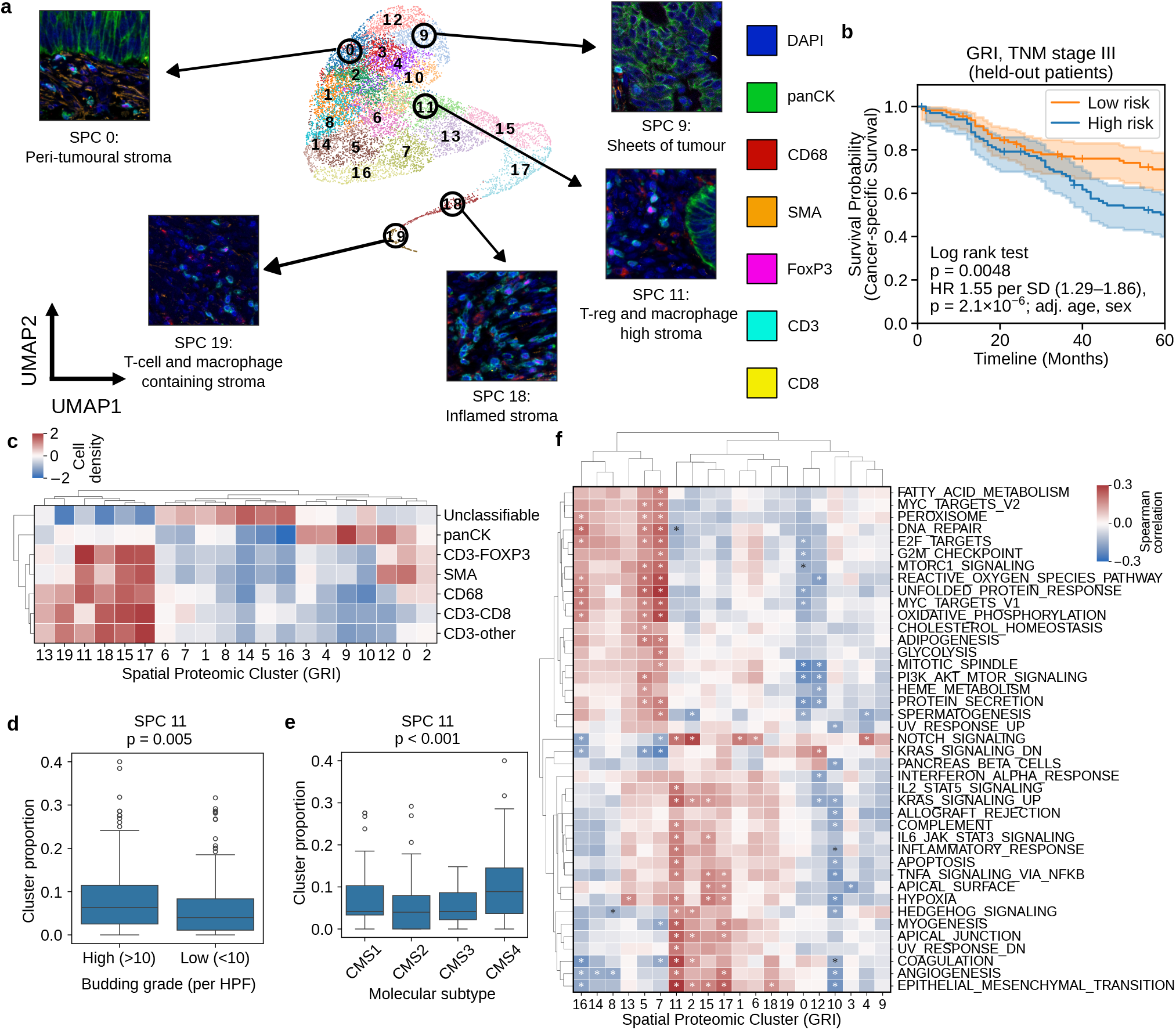
HyPERSTAC stratifies stage III colorectal carcinoma patients and identifies an immunosuppressive T_reg_- and macrophage-rich stromal cluster. (a) UMAP of the GRI tile embeddings, coloured by SPC, with example tiles (111 *µ*m) and pathologist descriptions. (b) Cancer-specific survival of held-out stage III patients (*n* = 215) by the HyPERSTAC risk score of Figure 2b: the score separates patients within a single stage (log-rank *p* = 0.0048; HR 1.55 per s.d., 95% CI 1.29–1.86, adjusted for age and sex). Shading, 95% CI. (c) Mean density of each cell type per SPC (z-score): stromal SPCs differ in immune composition, and SPC 11 is rich in FOXP3^+^ regulatory T cells and macrophages and poor in CD8^+^ T cells. (d) SPC 11 score by tumour budding grade. High grade *n* = 148, low grade *n* = 358 (patients with budding grade, both folds). Statistical significance from a two-tailed Mann-Whitney-U test. (e) SPC 11 score by consensus molecular subtype. CMS1 *n* = 60, CMS2 *n* = 170, CMS3 *n* = 23, CMS4 *n* = 60. Statistical significance from the Kruskal-Wallis test. (f) Correlation matrix showing the cluster proportion score against the single sample gene set enrichment analysis score of hallmark pathways. Pathways with at least one significant relationship with a cluster are shown. Values are Spearman correlation coefficients; * *p <* 0.01 after correction for multiple testing applied with the Benjamini-Hochberg method.

We found that enrichment for SPC 11 was associated with the high stroma content consensus molecular subtype (CMS) of CRC, CMS4 [33] as well as high grade tumour budding (Figures 5d, e). This is corroborated through pathway analysis, where we found that the proportion of SPC 11 on the slide correlated with transcriptomic modules of stromal remodelling such as epithelial-mesenchymal transition (EMT), myogenesis and angiogenesis, as well as those of immunosuppression via IL6/STAT3 signalling (Figure 5f) that has been shown to suppress anti-tumour effectors and promote regulatory immune cell function including T_reg_ and to promote tumour cell growth and survival [34, 35]. These observations support the notion that STAT3 signalling may promote an immunosuppressive environment throughout the tumour, which permits the evolution to a highly infiltrative phenotype with high grade tumour budding.

While similar observations have been subsequently made [36], these were not apparent at the time of data-generation. This illustrates the potential for HyPERSTAC to surface real, independently reproducible biological discovery autonomously, without human supervision.

### 2.5 Apical GLB1 expression is associated with low-grade EGFR-driven LUAD

In order to integrate the information obtained by Hy-PERSTAC from both LATTICe-A panels, we next used non-negative matrix factorisation (NMF) to combine the SPCs from both panels into 12 components (Figure 6a). These 12 components incorporate the information obtained from all SPCs, and enable patients to be described in terms of combined features from each panel (Figure 6a, c; Supplementary Figures S10a, b). We performed an associative survival analysis to guide subsequent interrogation of components, finding components 6 and 11 to be associated with favourable prognosis (Supplementary Figure S10c, d). On visual inspection of tumours with a high component 6 score, we observed that many of these tumours were of lepidic or papillary growth pattern, which are lower grade LUAD phenotypes (Figure 6b, c). Interestingly, the defining spatial proteomic feature was epithelial GLB1 expression localising to the apical aspect of the cell (Figure 6b, d). HyPERSTAC allows for direct discovery of subtle image features such as the subcellular localisation of expressed proteins, which are not captured by standard per-cell intensity features, and only partially by a bespoke segmentation-based polarity measurement that was designed post-hoc (Figure 6i, Supplementary Figure S12a).

**Figure 6.**
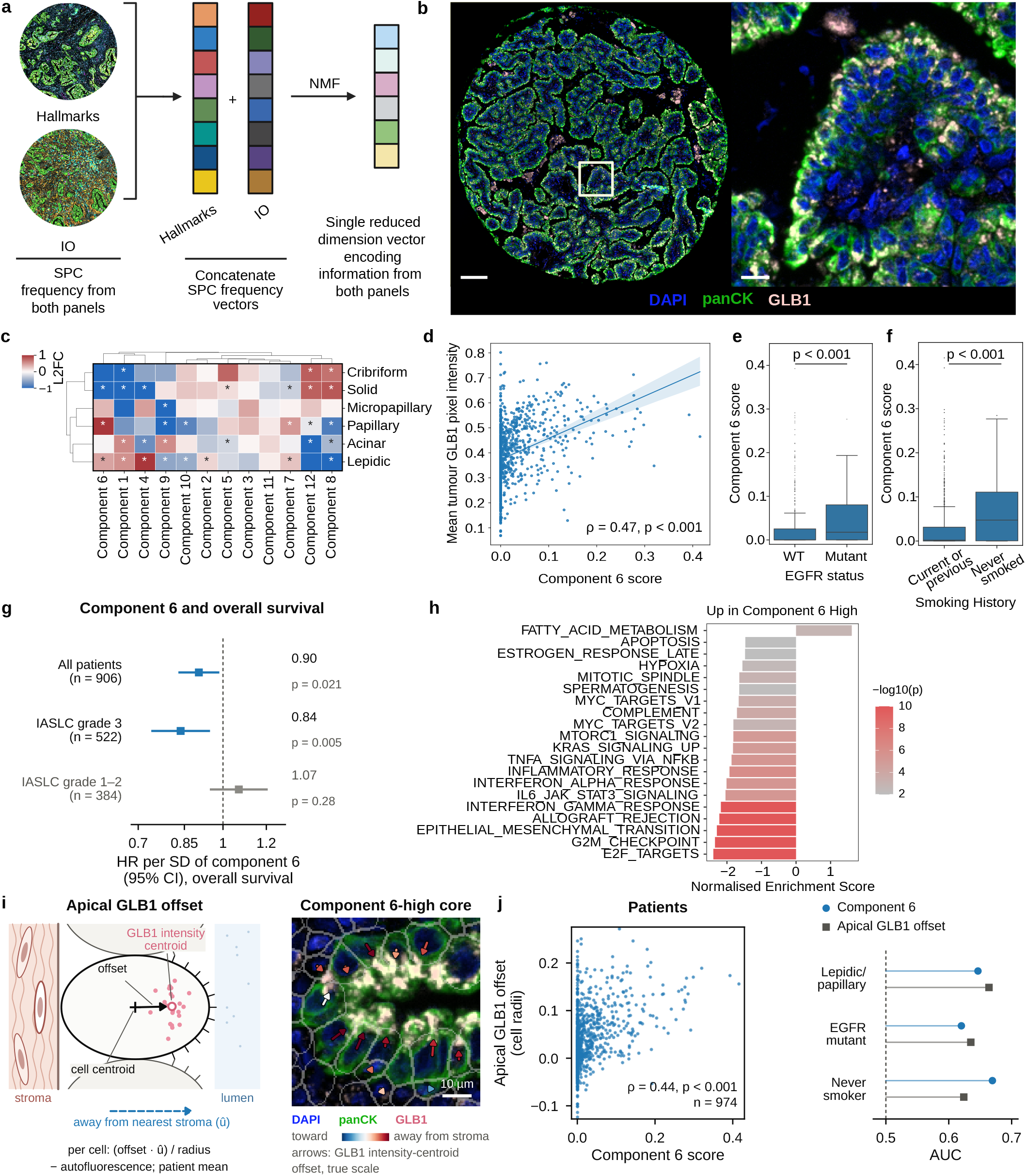
Cross-panel integration reveals a subcellular GLB1 pattern linked to EGFR-driven, low-grade lung adenocarcinoma. (a) Schematic of the integration of the two panels. SPC frequency vectors from the two panels are concatenated. A non-negative matrix factorisation model gives 12 components. (b) mIF image of a high component 6 core of LUAD with papillary architecture, displaying apical GLB1 intensity. Scale bars: core image 100 *µ*m; insert 10 *µ*m. (c) Component scores by predominant growth pattern (log_2_ fold change vs all cores; * adjusted *p <* 0.01): component 6 is enriched in low-grade lepidic and papillary tumours. A high score indicates a strong contribution of that SPC to that component’s score. (d) Component 6 rises only moderately with mean tumour-cell GLB1 per patient (Spearman *ρ* = 0.47), as it reflects where GLB1 is located rather than how much there is. (e, f) Component 6 is higher in *EGFR*-mutant tumours (e; *n* = 638) and never-smokers (f; *n* = 801; both *p <* 0.001). (g) HR per s.d. of component 6 for overall survival: protective overall (*n* = 906), strongest in IASLC grade 3 (*n* = 522) and absent in grade 1–2 (*n* = 384). (h) Gene sets differing between component 6-high and -low tumours: high scores go with lower proliferation, EMT and inflammatory signalling. (i) The pattern made explicit. Left, the per-cell GLB1 off-set from the cell centroid is projected away from the nearest stromal cell and corrected for autofluorescence. Right, a component 6-high core (colours as in b; grey, cell outlines) with each offset indicated by an arrow (red, away from stroma): most point away from stroma. Scale bar, 10 *µ*m. (j) This explicit offset correlates with component 6 per patient (left; *ρ* = 0.44, *n* = 974) and reproduces its associations with growth pattern, *EGFR* and smoking (right; AUC).

In addition to a relationship with low grade morphology, component 6 was also associated with *EGFR* mutation and never-smoking (Figure 6e, f). This is a recognised phenotype in LUAD, with poorly understood biology, seen classically in never-smokers, women and patients of East Asian ancestry [37]. This observation gives us a glimpse into the biology of this phenomenon, and allows us to follow this up with focused functional experiments to understand its significance.

Having discovered the relationship between component 6 and well-differentiated morphologies, we went back to the survival associations. Higher component 6 scores were associated with longer overall survival across all patients (HR 0.90 per s.d., 95% CI 0.83– 0.98, p = 0.021; Figure 6g). Given the relationship with low grade tumours, we performed a sub-group analysis and found that this association was stronger in IASLC grade 3 tumours (HR 0.84, 95% CI 0.74– 0.95, p = 0.005) and absent in grade 1–2 tumours (HR 1.07, 95% CI 0.95–1.21, p = 0.28; Figure 6g).

This raises the possibility that this pattern of GLB1 expression is a marker of reduced biological virulence across the whole tumour, its presence in one area indicating that conventionally high-grade appearances elsewhere might actually be less deadly than expected from H&E morphology alone. This idea is further supported at the transcriptomic level, where component 6-high tumours showed downregulation of protumourigenic pathways such as those relating to proliferation and EMT (Figure 6h).

Because HyPERSTAC identified the apical localisation of GLB1 without it being specified in advance, we next asked whether it could be defined as an explicit measurement on segmented cells. For each GLB1^+^ tumour cell within 15 *µ*m of stroma, we measured the offset of the GLB1 intensity-weighted centroid from the cell centroid in the direction away from the nearest stromal cell, corrected for autofluorescence, and averaged it per patient (the apical GLB1 offset; Figure 6i, Supplementary Figure S12a). The apical GLB1 offset correlated with component 6 (Spearman *ρ* = 0.44, p = 3 *×* 10^*−*47^, *n* = 974) and discriminated lepidic/papillary-predominant growth (AUC 0.66, vs 0.65 for component 6), *EGFR* mutation (0.64 vs 0.62) and never-smoking status (0.62 vs 0.67) to a similar degree (Figure 6j). Together with the other explicit GLB1 features, it accounted for the associations of component 6 with *EGFR* mutation, never-smoking and overall survival (component 6 given the explicit features, p = 0.87 for overall survival), whereas component 6 retained its association with lepidic/papillary growth (odds ratio 1.49 per s.d., p = 1.5 *×* 10^*−*5^) and a residual trend in grade 3 tumours (p = 0.09; Supplementary Figure S12). HyPERSTAC therefore discovered a subcellular pattern that could then be captured by a bespoke segmentation feature, designed after the fact, providing a route from hypothesis-free discovery to a candidate assay.

This phenomenon is highly directional. In component 6-high patients, GLB1 in tumour cells was displaced away from stroma about three times more than in patients with a component 6 score of zero, at every distance from the nearest air space, whereas the size of the offset beyond the autofluorescence null was similar in both groups (about 0.2 cell radii; Supplementary Figure S13a). Orienting the measurement by the density of all nearby stromal cells instead of the nearest stromal cell gave the same feature, whereas definitions oriented towards the nearest lumen were less strongly associated with component 6, and no alternative improved the association with survival (Supplementary Figure S13b).

Why GLB1 should be displaced away from stroma, and why in these tumours, may be explained by their differentiation. Lepidic/papillary, *EGFR*-mutant adenocarcinomas of never-smokers are largely of the terminal respiratory unit type, which retains alveolar type II (AT2) cell differentiation [38]. AT2 cells store surfactant in lamellar bodies, lysosome-related organelles that contain lysosomal hydrolases including *β*-galactosidase [39, 40] and are delivered to the apical membrane for secretion [41], and epithelial cells orient their apical pole relative to the basement membrane [42]. Apical GLB1 may therefore mark retained AT2-type secretory polarity, which would explain why the direction of GLB1, rather than its amount, distinguished component 6-high tumours, and why that direction was set by the stroma rather than by a nearby air space. Consistent with this, in public data *GLB1* co-varied with an AT2/lamellar-body program in bulk tumours [32], at the protein level [43] and within malignant cells of a single-cell lung cancer atlas [44], independently of macrophage content, and was lowest in dedifferentiated (EMT and neuroendocrine) tumour-cell states (Supplementary Figure S13c,d). Furthermore, and in support of a more generalised intracellular polarity programme, in an independent spatial transcriptomic dataset [45], *GLB1 transcripts* were also displaced away from stroma within tumour cells (86^th^ percentile of 2,355 genes), and more so in AT2-like cells, as were transcripts of the lamellar-body transporter *ABCA3* (Supplementary Fig. S13e).

### 2.6 Spatial organisation of tumour infiltrating lymphocytes predicts prognosis in PD-L1^+^ patients

While the previous sections have focused on obtaining a global set of SPCs for each panel/cohort, HyPER-STAC can obtain more granular insights by examining SPCs discovered within targeted patient subsets. For example, we examined the morphological heterogeneity within only the PD-L1^+^ patients with a view to discovering subgroups with good or poor outcomes in order to improve future patient selection. Without retraining any models, we used HyPERSTAC’s representations to discover a new set of SPCs (Figure 7a, b), which we refer to as PD-L1^+^SPCs. We found that many of these PD-L1^+^SPCs highlighted recurrent patterns of tumour/immune interaction.

**Figure 7.**
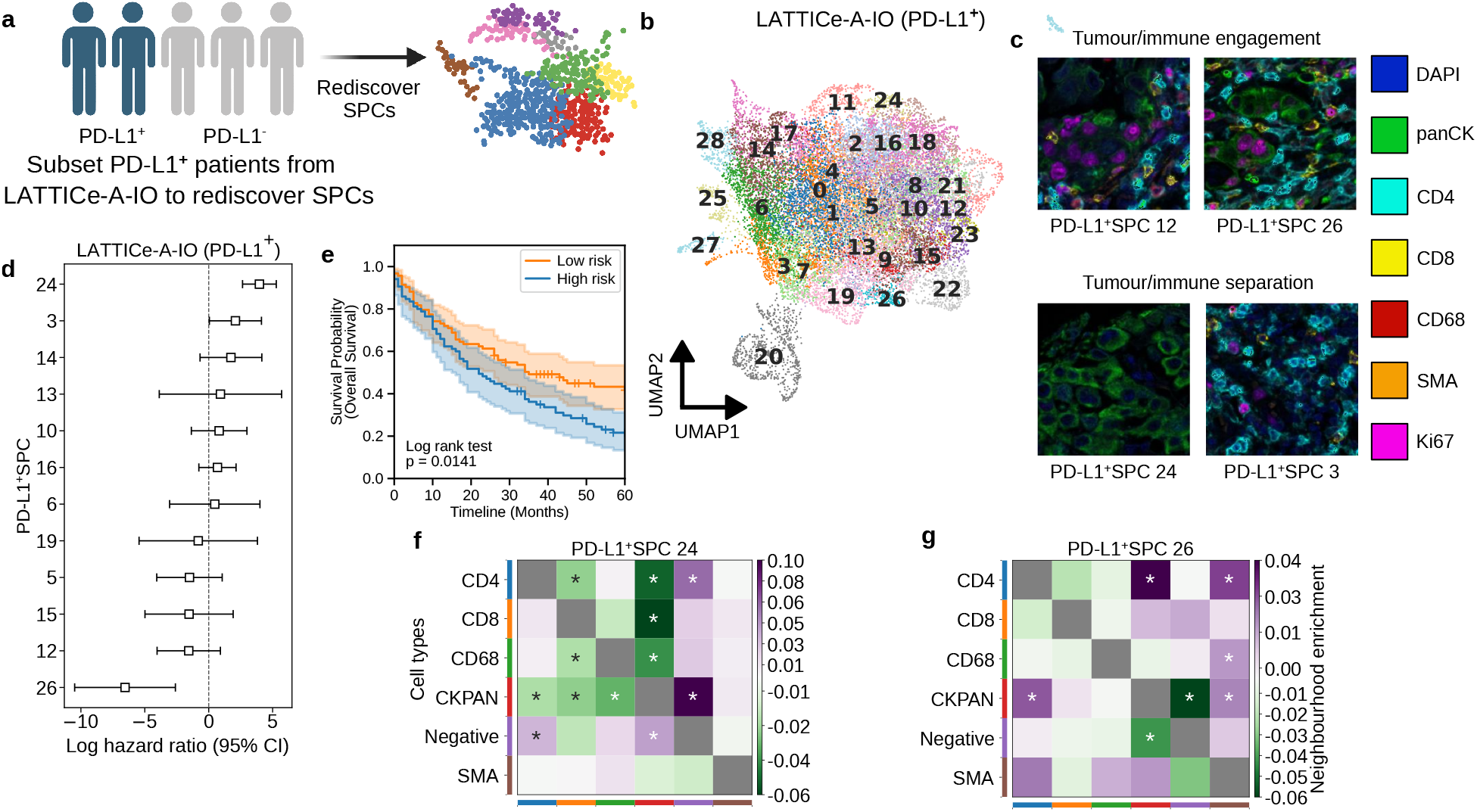
Spatial organisation of tumour-infiltrating lymphocytes predicts prognosis in PD-L1^+^ lung adenocarcinoma. (a, b) SPCs re-derived from PD-L1^+^ patients only (LATTICe-A-IO), without retraining the encoder (a), and their UMAP (b). (c) Example tiles (111 *µ*m): tumour and immune cells co-localise in PD-L1^+^SPCs 12 and 26 and are segregated in 24 and 3. (d) Log HR (95% CI) for overall survival of each PD-L1^+^SPC (fold-0 training patients, *n* = 101): separation-type SPC 24 is adverse and engagement-type SPC 26 protective. (e) Held-out PD-L1^+^ patients (*n* = 178) split by risk from these SPCs (log-rank *p* = 0.014): they stratify a group that PD-L1 status treats as uniform. Shading, 95% CI. (f, g) Difference in cell-type neighbourhood enrichment between cores high and low in PD-L1^+^SPC 24 (f) or 26 (g)(high minus low; purple, more contact in SPC-high cores; green, less; * *p <* 0.05). Where separation-type SPC 24 is frequent, CD4^+^ and CD8^+^ T cells and macrophages are depleted next to tumour cells; where engagement-type SPC 26 is frequent, CD4^+^ T cells are enriched next to tumour cells: the tile-level patterns in (c) extend to whole cores.

For example PD-L1^+^SPCs 24 and 3 show mutual exclusivity of tumour and immune cells, whereas in PD-L1^+^SPCs 12 and 26 they tend to co-localise within the 111 *µ*m tile region (Figure 7c). A cox model of these PD-L1^+^SPC frequencies stratified held-out PD-L1^+^ patients into risk groups (log-rank *p* = 0.014; Figure 7d, e).

We quantified these spatial relationships by carrying out a neighbourhood analysis, measuring the differential enrichment of immune cells near tumour cells between patients enriched or depleted in the target PD-L1^+^SPC. In cores from patients high in PD-L1^+^SPC 24, CD4^+^ and CD8^+^ T-cells and macrophages were all significantly depleted near tumour cells (Figure 7f), whereas in cores high in PD-L1^+^SPC 26, CD4^+^ T-cells were significantly enriched near tumour cells (Figure 7g). We found that the frequency of these co-localisation events were key drivers of prognosis in the PD-L1^+^ subset (Supplementary Figures S14a-d).

We also quantified the relationship between PD-L1^+^SPCs and the original set. We found that despite near equivalence between SPC 17 and PD-L1^+^SPC 26, this SPC was only predictive of prognosis in the PD-L1^+^ patient group (Supplementary Figure S14, Figure 7d).

These findings highlight the importance of context for the immune response to exhibit successful anti-tumour activity. The mere presence of tumour-infiltrating lymphocytes in PD-L1^+^ LUAD patients is insufficient to predict a good prognosis, and spatial measures are perhaps generally required to do so. To extend the idea further, it seems likely that a spatial measure of T-cell engagement may also prove valuable in predicting patient response to anti-PDL1 immunotherapy [46, 47].

## 3 Discussion

HyPERSTAC simplifies and democratises the interpretation of multiplex imaging by introducing unsupervised, image-first analysis to segmentation-based quantification as a powerful hypothesis-generating engine. We shift the focus of mIF analysis from intractable whole core images to SPCs, recurrent morphological and spatial proteomic patterns in LUAD and CRC which encode the spatial architecture of each sample. By using SPCs in combination with segmentation-derived motifs, HyPERSTAC is unconstrained by the limitations of solely cell segmentation- and/or lineage-based analyses which, when used alone, can be reductionist and prone to losing subtle morphological cues such as patterns of subcellular staining or extracellular detail. This permits important phenomena to be captured by consideration of the same image from different perspectives.

Crucially, SPCs are immediately interpretable and explainable, avoiding the *black-box* nature of many other AI methods [48]. HyPERSTAC builds on rather than replaces segmentation-based analysis: adding SPCs from any of four SSL methods or two foundation models improved the held-out prognostic performance of every established method we tested, including neighbourhood motifs, TRACERx-PHLEX, and MuSpAn, in all three datasets (Fig. 2). In our cohorts, joint-embedding SSL performed best: VICReg ranked highest overall, and VICReg and SimCLR were the only encoders whose SPCs, added to neighbourhood motifs, significantly improved on motifs alone in both LUAD panels, whereas MAE did so only in the IO panel (Supplementary Fig. S6). DINOv2, also a joint-embedding method, performed poorly in LATTICe-A, where its held-out embeddings collapsed onto a few clusters. Cohort-trained VICReg also outperformed the off-the-shelf multiplex foundation models KRONOS [30] and VirTues [25], implying that training on the cohort itself outperforms transfer from off-the-shelf foundation models. What distinguishes HyPERSTAC is less the choice of encoder than its integration with segmentation, which allows SPCs to be interpreted through, and evaluated alongside, the cell-level quantification that pathologists and established frameworks already use.

Some of these SPCs are candidate spatial biomarkers which are immediately translatable. For example, patient selection for immunotherapy is imperfect [1] and biomarkers beyond simple PD-L1 status have previously been explored [1, 49]. The density and spatial positioning of immune cells has been previously shown to be related to immunotherapy treatment response in LUAD [46, 47] and other solid malignancies [50]. Our findings support the notion that T-cell engagement is prognostically favourable for those with PD-L1^+^ tumours, and HyPERSTAC identified these recurrent patterns within our data without any supervision. Furthermore, we observed blunting of the successful anti-tumour immune response in regions showing high levels of tumour cell proliferation, as assessed by Ki67 expression. This could be due to impaired MHC I antigen presentation during mitosis as many normal cellular processes are disrupted, leading cells in-cycle to be relatively resistant to immune surveillance. Crucially, cell cycle genes are frequently upregulated in tumours resistant to immunotherapy, and direct targeting of the cell cycle holds promise in combination with immune checkpoint blockade [51]. While our cohorts lack treatment-paired immunotherapy response data, we believe our methodology to be useful for providing candidate spatial biomarkers both for immunotherapy or for novel therapeutic agents in clinical trials in the future. Furthermore, we expect that HyPERSTAC could be used to derive highly accurate spatial biomarkers from biologically focused single-plex immunohistochemistry, or oligo-plex (eg 3 or 4 coloured dyes) chromogenic assays, both of which are compatible with current diagnostic laboratory workflows, unlike mIF.

Other SPCs raise interesting avenues for further functional research. GLB1 is classically used as a protein marker of cellular senescence [52], but the punctate pattern of apical staining we observed in low-grade LUAD is distinct from that often seen in senescence, which appears much more diffuse [53, 54, 55, 56]. The prognostic implications of GLB1 in solid malignancies are also mixed [53, 54, 55, 56]. Despite no prior hypothesis about GLB1 polarity, once discovered the apical GLB1 pattern was useful as an explicit feature: when measured, directional subcellular GLB1 polarisation correlated with the original GLB1-associated cross-panel component (Spearman *ρ* = 0.44), reproduced most of its clinical associations and absorbed its association with overall survival in the whole cohort (Fig. 6i,j; Supplementary Fig. S12). This illustrates the pathway HyPERSTAC enables, from hypothesis-free discovery to a targeted, interpretable assay.

Two key limitations guide future work. First, while HyPERSTAC automates pattern discovery, mechanistic understanding requires integration with orthogonal, functional experimental data. Second, tile-based analysis captures local but not global tissue architecture; graph-based extensions or hierarchical transformers could help integrate spatial information from multiple scales.

As mIF and related spatial technologies mature, HyPERSTAC offers a framework to decode the *grammar* of tumour ecosystems, where spatial relationships between cells or intracellular protein distribution may be as prognostic as molecular alterations. By transforming images into hypothesis-generating engines, HyPERSTAC has the potential to greatly accelerate the discovery of next-generation biomarkers that integrate morphology, multiplex protein data, and cellular interactions, as well as driving biological discovery. Ultimately, such tools may shift pathology from descriptive reporting to predictive ecosystem modelling, ushering in an era of truly spatial precision medicine. We envision HyPERSTAC as the foundation for a new class of spatial biomarkers, where AI-driven discovery guides targeted clinical assays.

## 4 Methods

### 4.1 Datasets

We used 1 mm TMA tumour cores from two cancer cohorts: the Leicester Archival Thoracic Tumour Investigatory Cohort – Adenocarcinoma (LATTICe-A) lung cancer cohort, which features *>*2,500 cores from 1,025 lung adenocarcinoma patients, and 0.6 mm cores from the Glasgow Royal Infirmary (GRI) colorectal cancer cohort, comprising 787 colorectal cancer patients. After segmentation, quality control and linkage to outcome data, 1,003 LATTICe-A patients (IO panel 995 patients and 2,557 cores; Hallmarks panel 996 patients and 2,595 cores) and 563 GRI patients (933 cores; 1–3 cores per patient) were analysed.

The LATTICe-A cohort was originally constructed under REC 14/EM/1159 (East Midlands REC), and the ongoing management of the resource by the Greater Glasgow and Clyde Biorepository was approved under an amendment granted by the Leicester South REC. Ongoing use of the collection is now managed under REC 16/WS/0207.

The GRI cohort included 787 patients with stage I-III colorectal cancer (CRC) who underwent primary surgical resection at the Glasgow Royal Infirmary between 1997 and 2013. Patients were excluded if they received neoadjuvant therapy, had an emergency operation, or died within 30 days of surgery. Tumours were staged using the 5th Edition AJCC/UICC-TNM system and clinicopathological data were collected with a minimum 5-year follow-up post-resection. The study was approved by the West of Scotland Research Ethics Committee (REC 4: 22/ws/0207) and data are stored in the Greater Glasgow and Clyde Safe-haven (SH21ON012).

On LATTICe-A, we applied two distinct staining panels: the immune cell phenotyping panel *LATTICe-A-IO* and the hallmarks of cancer panel *LATTICe-A-Hallmarks*. LATTICe-A-IO uses markers DAPI, pan-CK, CD68, SMA, CD4, CD8, and Ki67. LATTICe-A-Hallmarks uses markers DAPI, pan-CK, N-cadherin, Ca9, cC3, VWF, Ki67, PD-L1, and GLB1. We stained the GRI cohort with another immuno-oncology panel, featuring DAPI, pan-CK, CD68, SMA, FoxP3, CD3, and CD8.

Datasets were split randomly into two folds on the patient level to avoid data leakage where cores from the same patient were assigned to different folds. Where applicable, e.g. for survival analysis, two-fold cross validation was used, and for analysis of clusters, the first fold was always used for training with the second fold kept as a holdout test set of similar size. Any findings which are associative across the whole cohort are clearly marked in the text.

### 4.2 Image Acquisition

The GRI panel was prepared and imaged as previously described [27].

#### 4.2.1 LATTICe-A-IO

A multiplex assay consisting of 6 markers was applied to twenty-three 4 *µ*m-thick FFPE sections from the LATTICe-A cohort using the Ventana Discovery Ultra autostainer (Roche Tissue Diagnostics, RUO Discovery Universal v21.00.0019). Discovery CC1

(Roche Tissue Diagnostics, 950-123) was applied to the section for 64 minutes at 95 °C for antigen retrieval. The antibodies were applied as follows:

CD68 (D4B9C) (Cell Signaling Technologies, 76437) at 1:200 for 32 minutes, followed by Discovery OmniMap anti-Rb HRP (Roche Tissue Diagnostics, 760-4311) for 16 minutes, detected by Opal 690 (Akoya Bioscience, FP1497001KT) at 1:100 for 8 minutes; Pan-Cytokeratin (AE1/AE3) (Leica Biosystems AE1/AE3-601-L-CE) at 1:250 for 28 minutes, followed by Discovery OmniMap anti-Ms HRP (Roche Tissue Diagnostics, 05269652001) for 12 minutes, detected by Opal 620 (Akoya Bioscience, FP1495001KT) at 1:100 for 8 minutes; Smooth Muscle Actin (1A4) (Roche Tissue Diagnostics, 760-2833) for 32 minutes, followed by Discovery OmniMap anti-Ms HRP (Roche Tissue Diagnostics, 05269652001) for 12 minutes, detected by Opal 540 (Akoya Bioscience, FP1494001KT) at 1:200 for 8 minutes; CD4 (SP35) (Roche Tissue Diagnostics, 790-4423) for 1 hour, followed by Discovery OmniMap anti-Rb HRP (Roche Tissue Diagnostics, 760-4311) for 16 minutes, detected by Opal 570 (Akoya Bioscience, FP1488001KT) at 1:50 for 8 minutes; CD8*α* (C8/144B) (Cell Signaling Technologies, 70306) at 1:100 for 56 minutes, followed by Discovery OmniMap anti-Ms HRP (Roche Tissue Diagnostics, 05269652001) for 12 minutes, detected by Opal 520 (Akoya Bioscience, FP1487001KT) at 1:100 for 8 minutes; and Ki67 (30-9) (Roche Tissue Diagnostics, 790-4286) for 20 minutes, followed by Discovery OmniMap anti-Rb HRP (Roche Tissue Diagnostics, 760-4311) for 12 minutes, detected by Opal 650 (Akoya Bioscience, FP1496001KT) at 1:400 for 8 minutes. DAPI was applied as a nuclear counterstain (Roche Tissue Diagnostics , 05268826001). Whole slide images were acquired at 20x magnification using the PhenoImager HT (Akoya Bioscience, v1.0.13) and unmixed using Phenochart (Akoya Bioscience, v1.1).

#### 4.2.2 LATTICe-A-Hallmarks

For LATTICe-A-Hallmarks, 23 tissue microarrays containing three 1.0mm cores from 1025 adenocarcinoma lung cancer patients (3075 cores in total) were cut at 4 *µ*m on TOMO slides (Matsunami, ref. TOM-11) and baked at 60°C for 60 minutes. The staining was performed using the Ventana Discovery Ultra, Roche Tissue Diagnostics, v21.00.0019. Slides were dewaxed and retrieved using Discovery CC1 (Roche Tissue Diagnostics, ref. 06414575001) for 32 minutes at 95°C. A stripping step was performed after each opal using CC2 (Roche Tissue Diagnostics, ref. 05279798001) at 100°C.

The antibodies were applied in the following order:

- Cleaved Caspase 3 (Asp175), (Cell Signaling Technology, ref. 9661) at 1:200 for 32 minutes followed by Omnimap anti-rabbit HRP (Roche Tissue Diagnostics, ref. 05269679001) for 12 minutes, detected by Opal 650 (Akoya Biosciences, ref. FP1496001KT) at 1:200.
- N Cadherin [EPR1791-4], (Abcam, ref. 76011) at 1:50 for 64 minutes, followed by Omnimap anti-rabbit HRP (Roche Tissue Diagnostics, ref. 05269679001) for 32 minutes, detected by Opal 520, (Akoya Biosciences, ref. FP1487001KT) at 1:100.
- GLB1/Beta-galactosidase, (Abcam, ref. 96239) at 1:150 for 32 minutes, followed by OmniMap anti-rabbit HRP (Roche Tissue Diagnostics, ref. 05269679001) for 16 minutes, detected by Opal 540 (Akoya Biosciences, ref. FP1494001KT) at 1:200.
- Von Willebrand Factor [36B11] (Leica Biosystems, ref. NCL-L-vWF) at 1:25 for 32 minutes followed by Omnimap anti-mouse HRP, (Roche Tissue Diagnostics, ref. 05269652001) for 12 minutes, detected by Opal 570(Akoya biosciences, ref. FP1488001KT) at 1:100.
- Carbonic Anhydrase 9, (Abcam, ref. 15086) at 1:250 for 32 minutes, followed by Omnimap anti-rabbit HRP (Roche Tissue Diagnostics, ref. 05269679001) for 12 minutes, detected by Opal 690 (Akoya Biosciences, ref. FP1497001KT) at 1:200.
- Pan-Cytokeratin [AE1/AE3], (Leica Biosystems, ref. AE1/AE3-601-L-CE) at 1:250 for 32 minutes, followed by Omnimap anti-mouse HRP, (Roche Tissue Diagnostics, ref. 05269652001) for 12 minutes, detected by Opal 620 (Akoya Biosciences, ref. FP1495001KT) at 1:100.
- PD-L1 [E1L3N] (Cell Signalling Technology, ref. 13684) at 1:50 for 120 minutes followed by Ultramap anti-rabbit HRP (Roche Tissue Diagnostics, ref. 05269717007), detected by Opal 480 (Akoya Biosciences, ref. FP1500001KT) at 1:50.
- KI67 [30-9] (Roche Tissue Diagnostic, ref. 05278384001) for 20 minutes, followed by Omnimap anti-rabbit HRP (Roche Tissue Diagnostics, ref. 05269679001) for 12 minutes, detected by TSA-DIG (Akoya Biosciences, ref. FP1501001KT) for 24 minutes and Opal 780 (Akoya Biosciences, ref. FP1501001KT) for 60 minutes.
- QD DAPI (Roche Tissue Diagnostics, ref. 05268826001) was applied for 24 minutes as counterstain.

All antibodies were previously validated in a chromogenic DAB assay using the Ventana Discovery Ultra. A single staining slide was created for each antibody – fluorophore combination to create a spectral library in InForm (Akoya Biosciences, v2.4.2) and an unstained section was used as autofluorescence reference. Slides were mounted and whole slide images were collected using PhenoImagerHT (Akoya Biosciences, v1.0.13) at 10x magnification, TMA maps were applied using Phenochart (v1.1.0) and individual core images were acquired at 20x magnification, and all images were spectrally unmixed using a project-specific spectral library using InForm.

### 4.3 HyPERSTAC SSL Architecture and Training

Every iteration, tiles were distinctly augmented to create two views of the same image, and each augmented tile was mapped to a latent representation by an eight-channel ResNet-50 [57]. To accommodate the increased number of imaging channels compared to standard RGB images, the augmentation regime used was: random left/right/up/down flipping (*p* = 1), random cropping (scale between 0.75 and 1, *p* = 1), random augmentation of the background (addition of uniform values to all 0-valued pixels, *p* = 0.3), random rotation (*p* = 0.4), solarisation (*p* = 0.3), and gaussian noise (*p* = 1). The VICReg [17] loss was used. This is a non-contrastive loss that does not require negative samples to avoid dimensional collapse, is computationally efficient, and has shown good results in other medical imaging settings [58]. In comparison to generative models that fixate on pixel-level reconstruction [59, 13, 23], joint-embedding architectures such as VICReg are designed to capture more meaningful, abstract features while ignoring more redundant information such as their exact appearance or location. The loss consists of three terms: variance *v*(***z***_*i*_), invariance *s*(***z***_1_, ***z***_2_), and covariance *c*(***z***_*i*_), for embeddings ***z***_*i*_ *∈* R^*D*^, *i* = 1, 2. The variance and covariance terms regularise the embedding to avoid collapse to a trivial solution, either by predicting the same score for all inputs or predicting the same value for each element of the embedding, respectively. The invariance term minimises the difference between the embeddings of each augmented image, meaning that the model should only learn features which are invariant to the augmentation regime–those features which have semantic meaning.

For benchmarking, the SimCLR models were trained using the same implementation as the VICReg models, simply changing the loss function for InfoNCE [16] with a temperature of 0.5. For DINOv2 [28] and MAE [29], the original implementations were used; MAE was a ViT-B trained separately for each fold, analogous to the masked-autoencoder approach of CANVAS [13], and its data loader quantised the inputs to 8 bits. The multiplex-imaging foundation models KRONOS [30] and VirTues [25] were applied with their published pretrained weights. KRONOS was run with the marker identity of each channel supplied as input (a run without marker identifiers was used as a sensitivity analysis).

SSL models were trained for 100 epochs on an Nvidia H100 GPU with 80 Gb memory. VICReg and SimCLR were implemented in TensorFlow 2, using code from [60, 61, 62]. A batch size of 256 was used, and a warmup-cosine learning rate was used, with a maximum learning rate of 10^*−*4^ and a 10 epoch warmup schedule, starting from 0. A representation size of 2048 was used, with an embedding size of 8192 generated from a three layer projection head with learned batch normalisation and a ReLU activation at every layer, and a linear activation on the output. DiNOv2 and MAE used their original PyTorch implementations.

#### 4.3.1 Image Preprocessing

Images were tiled using a 224 *×* 224px sliding window approach (111 *µ*m at 0.497 *µ*m per pixel). Finding a tissue outline in mIF images is nontrivial, as there is a large amount of black space throughout the tissue in each image, so standard Otsu thresholding [63] segments out large portions of the tissue. To obtain the tissue mask, the DAPI channel was dilated with a kernel size of 50px, and the image was then downsampled by a factor of 10. A median filter was applied with a filter size of 20px, and the image was resized back to its native resolution. This created images with an approximate outline of the tissue based on the DAPI channel. Otsu thresholding was then used to obtain a tissue mask. These operations were carried out using scikit-image [64]. To prevent the model overfitting on the background noise or the shape of the tissue edge, tiles that were less than 50% tissue were excluded. Any core images containing less than 20 tiles were rejected due to the lack of sample size.

#### 4.3.2 Clustering and Analysis Methods

Representations of each tile in the train set were clustered using the Leiden community detection algorithm [65] with a resolution of 1. The learned mappings were then used to assign the test set to the same clusters. The frequency of each cluster label in all tiles across all cores from each patient was then used to construct a cluster frequency vector for that patient, normalised to 1. As the test set representations were mapped to the same clusters as the train set, this means each patient’s vector would have the same dimension. Where multiple panels were combined (LATTICe-A-IO and LATTICe-A-Hallmarks), we concatenated the cluster vectors and normalised the resulting vector to magnitude 1 for consistency.

The clusters were visualised using UMAP [66] projections into two dimensions to assess their distributions and check for any evidence of issues with the representation, cluster imbalance, or patient specificity. This was quantified in Figure S15. The distribution of markers across clusters was visualised to ensure there was no overfitting to marker intensity, as illustrated in Figures 3 and S3. Marker intensities were scaled to the range 0-1 for visualisation purposes.

### 4.4 HyPERSTAC Segmentation and Neighbourhood Analysis

#### 4.4.1 Preprocessing

Images were imported into Visiopharm (version 2021.09.2.10918). A deep learning analysis protocol package (APP), using the U-Net architecture [67], was trained over 288,000 iterations using 106 images with varied tissue morphologies as the training dataset. The APP utilised DAPI, autofluorescence, and cytokeratin channels for training to classify regions of necrosis, tumour, stroma, and background tissue in all LUAD core images to prepare for cell-level analysis.

For cell-level analysis, a second, custom deep learning APP based on the U-Net architecture was developed using annotated images from 15 LUAD cores, selected to capture a broad range of cell types and sizes. Manual annotations were used to generate background labels, with additional boundary labels assigned to each nucleus for training purposes. The APP was trained over 127,000 iterations using DAPI and Ki67 channels as inputs. Cytoplasmic labels were generated by nuclear expansion by 20 px (9 *µ*m) for tumour cells and 10 px (4.5 *µ*m) for stromal cells. Cells were then delineated by using the boundary probability heatmap. For each segmented cell, the spatial coordinates and the mean intensity of each marker in each cellular compartment were exported for downstream analysis.

All fluorescence intensities were log_10_(*x* + 1)-transformed and batch-corrected with ComBat [68], with the tissue microarray as the batch, as implemented in scanpy v1.10.1 [69].

For LATTICe-A-IO, mean whole cell intensity was used for all markers, except Ki67 where the mean nuclear intensity was used. Cells were phenotyped using k-means clustering, as implemented in scikit learn v1.4.0 [70] using *k* = 10. Where clusters were annotating the same cell type, these were merged to produce the final classification. For LATTICe-A-Hallmarks, the neighbourhood motifs (below) were obtained by k-means clustering of the unscaled mean marker intensities of each cell’s 50 *µ*m neighbourhood. Lineage calls for this panel, used for TRACERx-PHLEX and MuSpAn, are described in Section 4.7.

For GRI, cell phenotyping was performed using Visiopharm image analysis software. The ready-to-use app Phenoplex was applied, with the thresholds used to assign marker positivity described in Supplementary Table S1. Cells which had positivity for conflicting markers (eg. CD68 and SMA), as well as cells which were not positive for any of the markers were considered unclassifiable.

Cluster labels were projected onto original images and verified by a pathologist in all cases.

#### 4.4.2 Cell Quantification

Cells were assigned to tiles based on the centroid of their nuclei, inclusive of those cells which overlap tile boundaries. Measures of cell density per cluster were derived from cell density per tile and the mean taken of all tiles in the cluster. A similar approach was taken to mean marker intensity. A sample of 100 tiles from the test set in each cluster was also manually inspected by a senior pathologist, and textual descriptions were assigned to each cluster based on their marker enrichment and visual appearance. These descriptions aimed to be holistic, and capture the recurring patterns among the sampled tiles, which were largely consistent in visual appearance.

#### 4.4.3 Cell Neighbourhood Motif Discovery

For each cell, a local spatial neighbourhood was defined by a 50*µ*m radius boundary using a fast nearest-neighbour search. Phenotypic proportions within each neighbourhood were then calculated, and neighbourhoods containing 10 or fewer cells were excluded from downstream analysis.

To evaluate neighbourhood reproducibility, the same train/test splits were used as for SSL training. A mini-batch K-means algorithm was independently applied to each set using the neighbourhood phenotypic proportions with *k* set to identify the same number of clusters as the SSL algorithm. Each cohort’s model was cross-predicted on the opposing dataset to evaluate cluster stability. Single-cell cluster assignments were aggregated to output patient-level neighbourhood percentage compositions.

For LATTICe-A-Hallmarks, the motifs were instead obtained by k-means clustering of the unscaled mean intensities of the eight marker channels over each cell’s 50 *µ*m neighbourhood. These intensity-based motifs are the Hallmarks segmentation baseline in Fig. 2.

### 4.5 Data Analysis Methods

#### 4.5.1 LATTICe-A Cohort Integration

To integrate the information from both LATTICe-A panels, the cluster sets from each panel were merged using non-negative matrix factorisation [71] with *k* = 12. This yielded a set of twelve components which contained combined information from both panels.

#### 4.5.2 PD-L1^+^ Sub-Analysis

For the PD-L1^+^ patient cohort sub-analysis, patients were selected based on a tumour proportion score of 1% *≥* [72]. Clustering was then performed on the train set tiles from these patients, with tiles from PD-L1 negative patients excluded. These clusters were distinct from those obtained from the whole cohort, and were used to more closely interrogate the tissue architecture of this patient subset. A map illustrating the relationship between the clusters in each cohort is shown in Supplementary Figure S14. The PD-L1^+^SPC survival model used the same two patient-level folds and Kaplan–Meier protocol as the main analyses (risk groups of held-out patients split at the median predicted risk of the training fold, folds pooled), but with a Cox penalty of 0.05 rather than the penalty of 1 used for the benchmark analyses.

Cell neighbourhood analysis was performed using the Squidpy [73] and CellCharter [74] packages in Python. Each cell was assigned a label based on its lineage, and a directed adjacency matrix *A* is constructed by finding the 6 nearest spatial neighbours of each cell on a core-by-core basis. These adjacency matrices were used to calculate the asymmetric neighbourhood enrichment of each combination of cell types, which is the difference between observed and expected numbers of edges between cell types. Differential neighbourhood enrichment is the difference between the neighbourhood enrichment matrices in two conditions. The *p* value is calculated by randomly sampling by condition label for each sample with replacement. The *p* value is the proportion of cases where the magnitude of the permuted value is greater than the magnitude of the observed value. We sort patients by the prevalence of the PD-L1^+^SPC of interest, splitting them into two groups based on the median value (where the PD-L1^+^SPC is present in less than half of patients, the groups are presence of the PD-L1^+^SPC against no presence). We calculate the differential neighbourhood enrichment between patients in these groups for each cell type.

Co-localisation scores are calculated as the directed neighbourhood enrichment count between the two cell types. We consider scores in both directions, and a combined score, which is the sum of the enrichments in each direction (Supplementary Figure S14). These scores are used to separate patients into high- and low-enrichment groups, whose survival probabilities are presented.

#### 4.5.3 Survival Analysis

All survival analyses of feature sets (Fig. 2 and the associated Supplementary Figures) used one row per patient: the SPC labels of all tiles and the segmentation-derived features of all cores from a patient were pooled into a single patient-level vector. Models were evaluated by two-fold patient-level cross-validation using the folds defined above: each model was trained on one set and tested on the other, and the held-out C-index is the mean over the two folds. Overall survival in LATTICe-A was truncated at 5 years (1,825 days) and cancer-specific survival in GRI at 60 months, owing to the paucity of events beyond these times.

Cox proportional hazards models were fitted with lifelines [75] (CoxPHFitter) with an L2 penalty (penalizer) of 1 and robust (sandwich) variance estimates. A sensitivity analysis with a penalty of 0.05 gave the same conclusions. To control for the large number of variables relative to the sample size, features were selected using two-fold cross validation on the train set only. The training patients were split into two halves (alternate rows). For each feature, a univariate Cox model with the same penalty was fitted on one half and its C-index computed on the other, in both directions. Features with a mean C-index greater than 0.5 over the two directions were retained.

Within each panel, all benchmark models were evaluated on the same patients: those with outcome data and segmentation-derived neighbourhood motifs (LATTICe-A-IO 995, LATTICe-A-Hallmarks 996, GRI 563). When a patient lacked another feature block (for example, no tiles passed quality control for the SSL encoder: 6 patients in each LATTICe-A panel and 43 in GRI) or a statistic was undefined, the missing values were imputed with the training-fold median, and a binary missingness indicator was added for each distinct missingness pattern within the block. No patient was therefore dropped from any arm, and all arms were compared on identical patients.

Held-out discrimination was measured by Harrell’s C-index [76]. Differences in held-out C-index between two feature sets (ΔC) were assessed by paired boot-strap: test patients were resampled with replacement within each fold (1,000 resamples), with the fitted models held fixed, and ΔC was averaged over the two folds in each resample. We report percentile 95% confidence intervals (CIs) and two-sided bootstrap *P* values (twice the smaller of the proportions of resampled ΔC*≤* 0 and *≥* 0). CIs for individual C-index values were obtained by the same resampling. For Kaplan–Meier analysis, the cut-point in each fold was the median of the model’s predicted expected survival time in that fold’s training patients, and it was applied to the fold’s held-out patients: held-out patients with predicted expected survival below the training-fold median (that is, predicted risk above it) were classed as high risk and the rest as low risk, so no held-out information was used to set the cut-point. The held-out patients of both folds were then pooled and the groups compared by log-rank test. Forest plots were produced showing the log hazard ratios with 95% confidence intervals (Supplementary Figure S4).

#### 4.5.4 Clinicopathological Models

To test whether HyPERSTAC is prognostic beyond routine clinicopathological variables, we used the held-out risk score of the Fig. 2 HyPERSTAC model (neighbourhood motifs plus SPCs): the partial hazard that each fold’s model assigned to its held-out patients, z-scored within the fold. The held-out patients of both folds were pooled and analysed with unpenalised Cox models stratified by fold. We report hazard ratios (HRs) per standard deviation of the score with Wald 95% CIs and *P* values, and likelihood-ratio tests for the added score.

In LATTICe-A, the clinical model contained pathological stage (AJCC/UICC 8th edition; I, II, III– IV [77]), IASLC grade, age (per 10 years) and sex. Overall survival was truncated at 5 years. IASLC grade was derived from the recorded growth-pattern proportions using the IASLC 2020 rule [78]. After normalising the pattern proportions to sum to one, tumours with *≥* 20% high-grade patterns (solid, micropapillary and cribriform combined) were grade 3. Of the remaining tumours, those whose combined acinar and papillary proportion exceeded the lepidic proportion were grade 2, and the rest were grade 1. Invasive mucinous adenocarcinomas were excluded, because the IASLC grading system does not apply to them.

In GRI, the base model contained TNM stage (I, II, III; no analysed patient had stage 0 disease), age and sex. Cancer-specific survival was truncated at 60 months. An extended model added tumour budding (high vs low) [79], Klintrup–Mäkinen inflammatory grade (high vs low) [80], tumour–stroma percentage (high vs low) [81], mismatch repair (MMR) status and differentiation (poor vs other). MMR status was coded as proficient, deficient, or unknown/not tested, with unknown kept as a separate level; a sensitivity analysis treated unknown as missing. Within-stage Kaplan–Meier curves used the full-cohort risk groups (training-fold-median split) restricted to that stage, so no cut-point was chosen within a stage.

The added discrimination over the clinical model was the difference in Harrell’s C-index of the linear predictor (ignoring the fold strata) between the clinical model and the clinical model plus the score. Its 95% CI came from 1,000 bootstrap resamples of patients in which both models were refitted (percentile CI), and Harrell’s optimism-corrected ΔC is also given [82].

### 4.6 Comparison with Established Spatial Analysis Frameworks

We benchmarked HyPERSTAC against two established frameworks for the spatial analysis of segmented multiplexed images, TRACERx-PHLEX [9] and MuSpAn [10]. All were computed from the same Visiopharm single-cell segmentations and phenotypes that were used for the neighbourhood motifs, with cell centroids in *µ*m. We did not use the deep-imcyto segmentation step of TRACERx-PHLEX, which is designed for imaging mass cytometry; our images are Opal multiplex immunofluorescence, and the Visiopharm segmentations were used for every method. The cell types were: CD4, CD8, CD68, pan-CK, SMA and unclassified cells (IO); CD3^+^CD8^+^, CD3^+^FOXP3^+^ and other CD3^+^ T cells, CD68, pan-CK, SMA and unclassifiable cells (GRI); and tumour, endothelial and stromal cells for Hallmarks, taken from the lineage calls described in Section 4.7. Every feature set was evaluated with the patient-level survival pipeline described above, on the same patients and folds. All parameters, feature summaries and model arms were specified before any survival result for the corresponding features was computed.

#### 4.6.1 TRACERx-PHLEX

The Spatial-PHLEX module of TRACERx-PHLEX was re-implemented outside its Nextflow/Singularity pipeline. The GPU (cuGraph) breadth-first search was replaced with a CPU (SciPy) breadth-first search, and the squidpy [73] nearest-neighbour graph with an equivalent scikit-learn [70] graph. On the PHLEX example data, the port reproduced the original code exactly: all 249 tumour-cluster polygons and 24,276 cell-to-cluster assignments were identical, as were chain lengths and barrier fractions (maximum absolute difference 0).

We followed the default clustered-barrier workflow, with parameters in *µ*m. Tumour cells were clustered by DBSCAN [83] (*ε* = 25 *µ*m, minimum one cell), and cluster outlines were defined by *α*-shapes (*α* = 0.05). Only tumour cells in clusters of at least 2,000 *µ*m^2^ were used as barrier targets. Barrier scores were computed on a symmetrised 10-nearest-neighbour cell graph (maximum path length 1,000) for the following triplets:

- IO: CD8*→*pan-CK via SMA; CD8*→*pan-CK via CD68; CD4*→*pan-CK via SMA;
- >GRI: CD8 *→* pan-CK via SMA; via CD68; via FOXP3^+^ T cells.

For each triplet we summarised the mean all-paths barrier fraction, the mean all-paths adjacent-barrier fraction and the median shortest-path length. The clustering features were tumour-cluster density, fraction of tumour cells clustered, median cluster area, fraction of CD8 cells inside tumour clusters, and median signed distance of CD8 cells to the cluster boundary. Together these give 14 features per patient. Core values were aggregated per patient, weighted by the number of contributing cells. Cores with no source cells or no clustered tumour gave missing values. A sensitivity analysis removed unclassified cells before running PHLEX.

There is no published PHLEX configuration for the Hallmarks panel, so we pre-specified an analogue with the same implementation and parameters, in which endothelial cells replaced CD8 T cells. Its triplets were endothelial *→* tumour via stroma, Ki67^+^ tumour *→* endothelial via stroma, and CA9^+^ tumour*→* endothelial via stroma; for the last two, all endothelial cells were targets. The same clustering features were computed, with endothelial cells in place of CD8 cells.

#### 4.6.2 MuSpAn

Spatial statistics were computed for each core with MuSpAn v1.2.5 [10], in *µ*m. Each core’s tissue boundary was estimated as an *α*-shape [84] with a maximum edge length of 50 *µ*m; for fragmented cores this was increased to 100 or 200 *µ*m or replaced by the convex hull. All statistics used this boundary for edge correction. A statistic was set to missing for a core when a population involved had fewer than five cells.

The base family comprised:

- cross pair-correlation functions for tumour– immune, tumour–stroma and immune– myeloid/stroma pairs (annulus width and step 10 *µ*m, maximum radius 100 *µ*m), summarised as means over 0–30 and 30–100 *µ*m;
- J-functions (50 realisations) at 10, 20 and 30 *µ*m. The extended families were:
- *Weighted pair-correlation functions* [85] (Hallmarks only, because the IO and GRI exports contain no per-cell intensities). These were computed for endothelial or stromal cells relative to tumour cells marked by CA9, Ki67 or PD-L1 intensity. Marks were log-transformed and rescaled to [0, 1] using fixed cohort-wide 1st and 99th percentiles, and the function was averaged over 0–30 *µ*m at marks *≥*0.8 and *≤*0.2.
- *Cellular neighbourhoods*: 3-hop neighbourhoods on a Delaunay network [86] (edges *≤* 30 *µ*m). Their square-root-transformed compositions were clustered into *k* = 8 neighbourhood types by k-means fitted on 300,000 cells sampled from training-fold patients only, and patients were described by neighbourhood-type proportions.
- *Topographical correlation maps* for the main tumour–immune pair (Hallmarks: tumour– endothelial), with radius 50 *µ*m, kernel radius 150 *µ*m, kernel *σ* 50 *µ*m and mesh 10 *µ*m. Each map was summarised by its mean and by the fractions of the map above +1 and below *−*1.
- *Quadrat correlation matrices* (100 *µ*m quadrats, minimum 15 observations, *α* = 0.05): standardised effect sizes for three pre-specified pairs. This family was excluded for GRI by a sparsity rule fixed before computation and applied to a pilot set of cores without outcome data: CD8 cells were too sparse for the main pair to be estimated in most GRI cores.
- *Persistent homology* : Vietoris–Rips filtrations of the tumour and CD8 (Hallmarks: endothelial) point clouds (maximum dimension 1, maximum distance 50 *µ*m, at most 10,000 points). Features were the number and total persistence of H_0_ and H_1_ features persisting more than 10 *µ*m, and the maximum H_1_ persistence.

For the Hallmarks weighted pair-correlation functions, the reference population was randomly thinned to at most 500 cells per core. Core values were aggregated per patient, weighted by the size of the reference population; neighbourhood proportions were pooled over cells. This gave 54 (IO), 50 (Hallmarks) and 56 (GRI) MuSpAn features per patient. In the dimension-matched variant, all MuSpAn features were reduced by principal component analysis (fitted on the training fold) to the number of SPCs.

#### 4.6.3 Combination with SPCs

SPCs were added to each method, giving the corresponding HyPERSTAC model, by concatenating the patient’s SPC frequencies (VICReg encoder, from the Leiden clustering of the corresponding fold) to that method’s features. The combined model was compared with the method alone by paired bootstrap of held-out C-index.

### 4.7 Hallmarks Cell Phenotyping

The Visiopharm export for the Hallmarks panel contains one mean intensity per cell for eight unlabelled marker channels (14.76 million cells, 2,624 cores, 1,008 patients). Channels were assigned to markers by Spearman correlation of core-level mean intensities with those from an independent Cellpose [87] segmentation of the same images. Each channel matched exactly one marker (Spearman *ρ* 0.94–0.98).

The Visiopharm Hallmarks segmentation has no cell phenotypes, so lineage calls were made for TRACERx-PHLEX and MuSpAn. Thresholds were fitted to log_10_ intensity on fold-0 training patients, without outcome data, and applied to all cells; thresholds fitted on fold 1 agreed for *≥* 99.2% of cells. Cells were called tumour when pan-CK exceeded the boundary of a two-component Gaussian mixture model (GMM). Pan-CK^*−*^ cells were called endothelial when VWF exceeded the upper boundary of a three-component GMM fitted to pan-CK^*−*^ cells, and stromal otherwise. Within the tumour and non-tumour lineages, Ki67^+^ and CA9^+^ cells were called from the upper component of a three-component GMM. The epithelial calls agreed with Visiopharm’s tissue-region classifier: 90% of tumour-region cells and 4% of stroma-region cells were called tumour.

### 4.8 Composition-matched Tiles and Recovery of SPCs from Segmentation Features

Each tile was described by (i) the fractions of the six Visiopharm cell types and cell density, or (ii) these features plus the fractions of tumour, stroma and necrosis regions, the neighbourhood-motif fractions and the Cellpose Ki67^+^ fraction. A histogram gradient-boosted classifier (scikit-learn HistGradientBoostingClassifier [70]; 300 iterations, early stopping, seed 0) was trained to predict each tile’s SPC from these features. It was trained on fold-0 training tiles (*n* = 72,852) and scored by balanced accuracy on held-out tiles (*n* = 68,851; 28 SPCs; chance 1/28). The classifier was repeated in fold 1 (27 SPCs; balanced accuracy 0.23 with features (i) and 0.29 with features (ii)). As a reference, a logistic regression was fitted in fold 0 to the same labels on the first 50 principal components of the SSL embedding.

To test whether the SPC information that can be recovered from segmentation is prognostic, we replaced each tile’s SPC label with the classifier’s prediction and passed the patient-level frequencies of the predicted SPCs through the unchanged survival pipeline, in both folds. Predictions for training-fold tiles were cross-fitted (five-fold GroupKFold by patient), so that no tile was labelled by a model trained on it. Held-out tiles were labelled by the classifier trained on all training tiles. The held-out C-index was compared by paired bootstrap with that of the true SPCs computed on the same QC-passed tiles, and with that of the segmentation features used directly (patient-mean composition and density).

For the matched tiles, the prognostic association of each SPC was summarised by the univariate Cox HR per standard deviation of patient-level SPC frequency in the fold-0 training patients (overall survival truncated at 1,825 days), with held-out patients used for replication. Candidate SPC pairs had similar mean cell-type compositions (total-variation distance *≤* 0.10) and opposite training-fold HRs. The pair SPC 19 vs SPC 13 (largest difference in log HR) was pre-specified. Other pairs, including SPC 19 vs SPC 4 shown in Fig. 2d, were selected post hoc using the held-out direction of effect, and are illustrative. Within a pair, tiles came from different patients and from the central half of each SPC in embedding space; they had *≥* 25 cells, near-identical composition and usually the same dominant motif. Composition-adjusted Cox models of each pair’s share were fitted on held-out patients as an exploratory analysis.

### 4.9 Explicit GLB1 Polarity Feature

To test whether the apical GLB1 pattern captured by NMF component 6 admits a segmentation-based analysis, we constructed explicit per-cell GLB1 features. These features were designed post hoc, after component 6 had been identified. Cells were segmented with Cellpose [87] on the Hallmarks tiles, with nuclear masks expanded by 6 pixels to give cell masks. Only cells with *≥* 99% of their pixels imaged were used. Tumour cells were those with mean pan-CK intensity above the trough of its bimodal distribution and an area of at least 50 pixels. This gave 4.4 million tumour cells from 2,616 cores; 989 patients had at least 50 tumour cells.

For each tumour cell, the GLB1 weight of each pixel was its intensity minus the cell’s 10th-percentile intensity, clipped at zero. The offset was the vector from the geometric cell centroid to the GLB1-intensity-weighted centroid. For GLB1^+^ tumour cells within 15 *µ*m of stroma, the offset was projected onto the unit vector pointing away from the nearest panCK^*−*^ cell. To remove artefacts of mask geometry, the same projection computed from the autofluorescence channel was subtracted. The result was divided by the equivalent cell radius 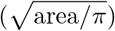 and averaged per patient over the qualifying cells. We also computed two measures of GLB1 texture: the share of a cell’s GLB1 signal held by its brightest 10% of pixels (higher for more concentrated staining), and granularity, the share of signal in bright puncta up to about 2 µm across (a white top-hat filter with a 1 µm-radius disk). We compared this definition with alternatives specified before they were computed: the direction away from a Gaussian-weighted density of all stromal (panCK^*−*^) cells within 3*σ* (*σ* = 10, 15 or 25 *µ*m; at least three stromal cells), the direction towards the nearest lumen (cell-free, low-autofluorescence space), and the bisector of the two. Each was compared with the nearest-stromal-cell definition by the difference in Spearman correlation with component 6 (paired bootstrap, 1,000 resamples of patients), and the nearest-stromal-cell definition was retained. In an exploratory analysis, GLB1^+^ tumour cells were binned by distance to the nearest lumen, and the autofluorescence-referenced projections towards the lumen and away from stroma, and the offset magnitude in excess of the autofluorescence offset, were averaged per patient and then across patients (boot-strap CIs over patients) for patients in the top quartile of component 6 and those with component 6 = 0.

*External expression and proteomic data*. To test whether GLB1 tracks the AT2 program, we defined an AT2/lamellar-body score as the mean *z*-score of *SFTPC, SFTPB, SFTPA1, SFTPA2, SFTPD, ABCA3, LAMP3, NAPSA, LPCAT1* and *SLC34A2*, and a macrophage score from *CD68, CD163, MARCO, C1QA, C1QB, MSR1, FABP4* and *MRC1*. We used TCGA-LUAD RNA-seq (516 primary tumours, one per patient; log_2_(TPM+1) from the GDC Xena hub) [32], CPTAC LUAD protein and RNA abundance (110 tumours; cBioPortal) [43], and the LuCA core single-cell atlas [44], from which malignant LUAD cells of primary tumours were averaged per patient (81 patients with *≥* 50 malignant cells; counts normalised to 10,000 per cell and log-transformed; genes *z*-scored within dataset). Associations were Spearman correlations, with partial rank correlations adjusting for the macrophage score and, in CPTAC, tumour purity; 95% CIs used Fisher’s *z* with standard error 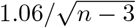.

*Subcellular spatial transcriptomics*. We used a public 10x Genomics Xenium Prime 5K dataset of one FFPE lung adenocarcinoma (Xenium Onboard Analysis 3.0.0) [45], whose panel includes *GLB1, ABCA3, LAMP1* and *LAMP3*. Cells were classified using the vendor’s graph-based clusters: clusters with a mean epithelial marker *z*-score (*EPCAM, CDH1, CLDN7, MUC1, EHF, GRHL2*) above 0.5 and above the fibroblast, endothelial and immune scores were epithelial, and all other cells were stromal. For epithelial cells (*≥* 30 transcripts) within 15 *µ*m of a stromal cell and each gene with at least two transcripts (quality value *≥* 20) in the cell, we projected the offset of the gene’s transcript centroid from the cell’s all-transcript centroid onto the direction away from the nearest stromal cell, in units of the equivalent cell radius; referencing to the all-transcript centroid plays the role of the autofluorescence reference in the mIF feature. Genes were summarised by their mean offset over cells (genes with *≥* 200 cells), and 95% CIs used a block bootstrap over 200 *µ*m tiles.

All tests used the Fig. 6 cohort: non-mucinous adenocarcinoma, overall survival not truncated, unpenalised models. Associations with component 6 were tested by Spearman correlation, and by partial correlation controlling for mean GLB1 and luminal-edge fraction. Associations with lepidic/papillary-predominant growth, EGFR mutation and never-smoking were tested by AUC (Mann–Whitney test, Benjamini–Hochberg adjusted [89]) and by multivariable logistic regression on component 6, mean GLB1, the apical offset, granularity and luminal-edge fraction. Associations with overall survival were tested by univariable Cox models (HR per standard deviation), overall and within IASLC grade 3, and by nested Cox models (Wald and likelihood-ratio tests).

### 4.10 Gene Expression and Genotyping

LATTICe-A gene expression was acquired and preprocessed as previously described [90]. For GRI, FFPE tissue digestion was carried out to obtain the tissue lysate and combined with detector oligos which annealed to the targeted RNA and ligated. Amplification of ligated oligos was then performed using a unique set of primers for each sample, introducing a sample-specific barcode and Illumina adaptors. Barcoded samples were pooled into a single library and run on an Illumina HiSeq 2500 High Output v4 flowcell. Sequencing reads were demultiplexed using BCL2FASTQ software (Illumina, USA). FASTQ files were aligned to the Human Whole Transcriptome v2.0 panel, which consists of 22,537 probes.

Differential expression analysis was performed in DESeq2 v1.40.2 [91], GSEA in fgsea v1.26.0 [92] and ssGSEA in GSVA v1.48.3 [93] using the MSigDb hallmarks database. TCGA-LUAD RNA-Seq was obtained from Xena Browser [94]. Cell deconvolution was done with MCP Counter v1.2.0 [31].

1 mm cores of tumour tissue were taken for DNA sequencing at the same time as cores were taken for TMA construction. Cores for sequencing were taken last to minimise DNA contamination from the previous case (ie typically after 12 passes of the coring needle through the case to be sequenced). DNA was extracted from tumour cores using the QIAmp FFPE DNA tissue kit (Qiagen) according to the manufacturer’s instructions. Recovered DNA was subsequently assessed for quantity and quality (DIN) using an Agilent Tapestation 4200 with Genomic screen tape. Where possible, 10 ng (but not less than 2 ng) of DNA was then sequenced using an ISO 15189 accredited custom Ampliseq based assay (MGP-4-DNA, Sarah Cannon Molecular Diagnostics). Library preparation was performed manually with templating and chip loading on an Ion ChefTM and sequencing on an Ion GeneStudioTM S5 Prime. Data analysis was undertaken using a hybrid bioinformatics pipeline comprising the native Torrent Suite (5.10.2) Variant Caller (5.10.1.20) combined with in-house developed applications for error checking, quality control, variant ‘binning’ and HGVS nomenclature construction. The assay was validated with a limit of detection for SNVs and small indels of 2.5% VAF in hotspot regions from *>*30 genes. The absolute VAF threshold for reporting variants as detected was adjusted upwards as required in many of the older specimens in order to allow for extensive ‘fixation’ associated background noise. The assay was also validated for the detection of copy number gains in 6 genes (EGFR, ERBB2, KRAS, KIT, MET & PIK3CA) in specimens with greater than 20% tumour cellularity. Samples which did not yield full panel coverage or which failed to meet other minimum quality control criteria were excluded.

TP53 status was taken from this targeted panel for LATTICe-A (641 patients with a result) and, for TCGA-LUAD, from the Xena whole-exome mutation calls (any non-silent TP53 variant; 495 patients) [94]. To test whether TP53 mutation explained the association between T-cell density, proliferation and survival (Supplementary Figure S9d, e), the T-cell effect was estimated within each Ki67 *×* TP53 group by univariable Cox regression of overall survival (truncated at 60 months), and Cox models with T-cell × Ki67 and T-cell *×* TP53 interaction terms were compared by AIC.

## Acknowledgements

The authors would like to extend their gratitude to: Catherine Winchester, CRUK Scotland Institute, for critically appraising this manuscript; Naveed Khan, CRUK Scotland Institute, for management of the high performance computing facility; the NHS Research Scotland (NRS) Greater Glasgow and Clyde Biorepository and Glasgow Tissue Research Facility for their ongoing management of the LATTICe-A and GRI cohorts; the CRUK Scotland Institute Deep Phenotyping Advanced Technology Core Facility (Research Resource Identifier [RRID]: SCR 027366) for the preparation of the images; the Academic Unit of Surgery at the Glasgow Royal Infirmary for the clinical database management; and Philip Bennett from Sarah Cannon Molecular Diagnostics, Part of HCA Healthcare UK for the preparation of the LATTICe-A genomic dataset.

## Author Contributions

LF and KR designed and executed the experiments, and prepared the manuscript. PH provided the GRI cohort images and contributed to their analysis. SS helped with the analysis of the LATTICe-A-Hallmarks images. FS assisted with the KRONOS foundation model benchmarking. LSS preprocessed and prepared the gene expression data for the GRI cohort. IRP analysed the neighbourhood motifs, designed the panels, and FB, RLB, SM, and LOJ performed the staining and imaging for the LATTICe-A images. DM, CRW, and MS constructed the clinicopathological and survival data for the LATTICe-A cohort. NM provided the histological assessment for the GRI cohort. CJM provided computational resources and feedback on experimental design. DC provided helpful feedback in the development of the method and production of the manuscript. JE and CR supervised the production of the GRI cohort. EWR supervised the experiments and provided feedback on experimental design and manuscript production. JLQ and KY designed the experiments, edited the manuscript, and jointly supervised the research.

## Data and Code Availability Statement

The codebase used to produce the results in this manuscript are available github.com/lucasfarndale/HyPERSTAC. Raw mIF images from the LATTICe-A cohort, with overall survival time and status, are available from the EBI BioImage Archive (S-BIAD3742); other clinical data are restricted (see below). Raw mIF images from the GRI cohort are available from JE upon reasonable request. Tile representations from both LATTICe-A and GRI are available on Zenodo (DOI: 10.5281/zenodo.17349843). Clinical data from the LATTICe-A cohort is restricted, and is held by NHS Greater Glasgow and Clyde Biorepository as custodians. Data access can be requested, and any such request will be reviewed and released under their research ethics committee-approved tissue bank protocols. Requests will be reviewed and approved within 6–8 weeks and will be accompanied by a data sharing agreement detailing the conditions and restrictions of use and publication.

## Funding Statement

LF is supported by the MRC, United Kingdom grant MR/W006804/1. KR is supported by a Jean Shanks Foundation and Pathological Society of Great Britain clinical PhD fellowship. The JLQ lab (SM, RLB, FB, IRP, JLQ) is funded by the Mazumdar-Shaw Chair Endowment and CRUK Scotland Institute core funding (A31287). LOJ is funded by the CRUK Scotland Centre CTRQQR-2021\100006. The JE lab (JE, PH,

LSS) is funded by CRUK (CTRQQR-202 \100006) and Innovate UK (42497). CJM acknowledges support a core programme award to CJM (A29801) and NC3Rs DA3RT grant (APP51238). EWR acknowledges CRUK core funding (A1920). EWR and KY are supported by funding from Prostate Cancer UK (MA-TIA22-001). KY acknowledges support from Cancer Research UK (EDDPGM-Nov21 100001 and DRCMDP-Nov23 100010), BBSRC BB V016067 1, and EU Horizon 2020 grant ID: 101016851. This work was supported by Cancer Research UK core funding to the Scotland Institute (A31287).

## Competing Interest Statement

DC, JLQ, and KY are co-founders and shareholders of TileBio Ltd.

## Notes

https://github.com/lucasfarndale/MultiplexSSL/

### Competing Interest Statement

David Chang, John Le Quesne, and Ke Yuan are co-founders and shareholders of TileBio Ltd.

### Summary of Updates

The method has been updated, with substantial new benchmarking and a dataset release

## References

[1] S. Lu et al. Comparison of Biomarker Modalities for Predicting Response to PD-1/PD-L1 Check-point Blockade: A Systematic Review and Meta-analysis. JAMA Oncology, 5(8):1195–1204, August 2019.

[2] A. T. Shaw et al. Crizotinib in ROS1-Rearranged Non–Small-Cell Lung Cancer. New England Journal of Medicine, 371(21):1963–1971, November 2014.

[3] M. Reck et al. Pembrolizumab versus Chemotherapy for PD-L1–Positive Non–Small-Cell Lung Cancer. New England Journal of Medicine, 375(19):1823–1833, November 2016.

[4] E. L. Kwak et al. Anaplastic Lymphoma Kinase Inhibition in Non–Small-Cell Lung Cancer. New England Journal of Medicine, 363(18), October 2010.

[5] H. Hampel et al. Screening for the Lynch Syndrome (Hereditary Nonpolyposis ColorectalCancer). New England Journal of Medicine, 352(18):1851–1860, May 2005.

[6] A. Passaro et al. Cancer biomarkers: Emerging trends and clinical implications for personalized treatment. Cell, 187(7):1617–1635, March 2024.Publisher: Elsevier.

[7] L. Keren et al. A structured tumor-immune microenvironment in triple negative breast cancer revealed by multiplexed ion beam imaging. Cell,174(6):1373–1387.e19, 2018.

[8] C. M. Schürch et al. Coordinated cellular neighborhoods orchestrate antitumoral immunity at the colorectal cancer invasive front. Cell,182(5):1341–1359.e19, 2020.

[9] A. Magness et al. Deep cell phenotyping and spatial analysis of multiplexed imaging with TRACERx-PHLEX. Nature Communications, 15(1):5135, 2024.

[10] J. A. Bull et al. MuSpAn: a toolbox for multi-scale spatial analysis. Nature Communications, 17, 2026.

[11] Y. Si et al. FICTURE: scalable segmentation-free analysis of submicron-resolution spatial transcriptomics. Nature Methods, 21(10):1843–1854, 2024.

[12] C. C. Liu et al. Robust phenotyping of highly multiplexed tissue imaging data using pixel-level clustering. Nature Communications, 14(1):4618, 2023.

[13] J. Tan et al. Characterization of tumour heterogeneity through segmentation-free representation learning on multiplexed imaging data. Nature Biomedical Engineering, pp. 1–15, 2025.

[14] D. Jiménez-Sánchez et al. Weakly supervised deep learning to predict recurrence in low-grade endometrial cancer from multiplexed immunofluorescence images. NPJ Digital Medicine, 6(1):48, 2023.

[15] M. Harary et al. Fluoroformer: Scaling multiple instance learning to multiplexed images via attention-based channel fusion. arXiv preprint arXiv:2411.08975, 2024.

[16] T. Chen et al. A simple framework for contrastive learning of visual representations. In International conference on machine learning, pp.1597–1607. PmLR, 2020.

[17] A. Bardes et al. Vicreg: Variance-invariance-covariance regularization for self-supervised learning. arXiv preprint arXiv:2105.04906, 2021.

[18] R. J. Chen et al. Towards a general-purpose foundation model for computational pathology. Nature Medicine, 30(3):850–862, March 2024. Publisher: Nature Publishing Group.

[19] X. Wang et al. Transformer-based unsupervised contrastive learning for histopathological image classification. Medical Image Analysis,81:102559, October 2022.

[20] E. Vorontsov et al. A foundation model for clinical-grade computational pathology and rare cancers detection. Nature Medicine, pp. 1–12,July 2024. Publisher: Nature Publishing Group.

[21] H. Xu et al. A whole-slide foundation model for digital pathology from real-world data. Nature, 630(8015):181–188, June 2024. Publisher: Nature Publishing Group.

[22] A. Claudio Quiros et al. Mapping the landscape of histomorphological cancer phenotypes using self-supervised learning on unannotated pathology slides. Nature Communications, 15(1):4596, 2024.

[23] E. Wu et al. Rosie: Ai generation of multiplex immunofluorescence staining from histopathology images. Nature Communications, 16(1):1–13, 2025.

[24] G. Atarsaikhan et al. Self-supervised learning enables unbiased patient characterization from multiplexed cancer tissue microscopy images.bioRxiv, pp. 2025–03, 2025.

[25] J. Wenckstern et al. Ai-powered virtual tissues from spatial proteomics for clinical diagnostics and biomedical discovery. arXiv preprint arXiv:2501.06039, 2025.

[26] D. A. Moore et al. In situ growth in early lung adenocarcinoma may represent precursor growth or invasive clone outgrowth—a clinically relevant distinction. Modern Pathology, 32(8):1095–1105, 2019.

[27] P. Hatthakarnkul et al. Protein expression of s100a2 reveals it association with patient prognosis and immune infiltration profile in colorectal cancer. Journal of Cancer, 14(10):1837, 2023.

[28] M. Oquab et al. Dinov2: Learning robust visual features without supervision. arXiv preprint arXiv:2304.07193, 2023.

[29] K. He et al. Masked autoencoders are scalable vision learners. In Proceedings of the IEEE/CVF conference on computer vision and pattern recognition, pp. 16000–16009, 2022.

[30] M. Shaban et al. A foundation model for spatial proteomics. arXiv preprint arXiv:2506.03373, 2025.

[31] E. Becht et al. Estimating the population abundance of tissue-infiltrating immune and stromal cell populations using gene expression. GenomeBiology, 17(1):218, October 2016.

[32] Cancer Genome Atlas Research Network. Comprehensive molecular profiling of lung adenocarcinoma. Nature, 511(7511):543–550, 2014.

[33] J. Guinney et al. The consensus molecular subtypes of colorectal cancer. Nature Medicine, 21(11):1350–1356, November 2015. Publisher:Nature Publishing Group.

[34] D. E. Johnson et al. Targeting the il-6/jak/stat3 signalling axis in cancer. Nature reviews Clinical oncology, 15(4):234–248, 2018.

[35] K. A. Pennel et al. Jak/stat3 represents a therapeutic target for colorectal cancer patients with stromal-rich tumors. Journal of Experimental & Clinical Cancer Research, 43(1):64, 2024.

[36] P. Hatthakarnkul et al. The characteristic of epithelial-specific phenotypes and immunosuppressive microenvironment in the context of tumour budding in colorectal cancer. bioRxiv, pp. 2025–03, 2025.

[37] H. Shigematsu et al. Clinical and biological features associated with epidermal growth factor receptor gene mutations in lung cancers. Journal of the National Cancer Institute, 97(5):339–346,2005.

[38] Y. Yatabe et al. EGFR mutation is specific for terminal respiratory unit type adenocarcinoma. The American Journal of Surgical Pathology, 29(5):633–639, 2005.

[39] G. E. Hook and L. B. Gilmore. Hydrolases of pulmonary lysosomes and lamellar bodies. Journal of Biological Chemistry, 257(15):9211–9220,1982.

[40] R. H. Glew et al. Sequential changes of lamellar body hydrolases during ozone-induced alveolar injury and repair. The American Journal of Pathology, 134(5):1143–1150, 1989.

[41] A. V. Andreeva et al. Regulation of surfactant secretion in alveolar type II cells. American Journal of Physiology. Lung Cellular and Molecular Physiology, 293(2):L259–L271, 2007.

[42] L. E. O’Brien et al. Rac1 orientates epithelial apical polarity through effects on basolateral laminin assembly. Nature Cell Biology, 3(9):831–838, 2001.

[43] M. A. Gillette et al. Proteogenomic characterization reveals therapeutic vulnerabilities in lung adenocarcinoma. Cell, 182(1):200–225.e35, 2020.

[44] S. Salcher et al. High-resolution single-cell atlas reveals diversity and plasticity of tissue-resident neutrophils in non-small cell lung cancer. CancerCell, 40(12):1503–1520.e8, 2022.

[45] 10x Genomics. Xenium Prime 5K In Situ Gene Expression with Cell Segmentation: human lung adenocarcinoma (FFPE). https://www.10xgenomics.com/datasets/xenium-human-lung-cancer-post-xenium-technote, 2024. Dataset Xe-nium Prime Human Lung Cancer FFPE, Xenium Onboard Analysis 3.0.0.

[46] F. Ghiringhelli et al. Immunoscore immune checkpoint using spatial quantitative analysis of CD8 and PD-l1 markers is predictive of the efficacy of anti-PD1/PD-l1 immunotherapy in non-small cell lung cancer. EBioMedicine, 92:104633, 2023.

[47] J. Monkman et al. Spatial insights into immunotherapy response in non-small cell lung cancer (NSCLC) by multiplexed tissue imaging.Journal of Translational Medicine, 22(1):239, 2024.

[48] C. Rudin. Stop explaining black box machine learning models for high stakes decisions and use interpretable models instead. Nature Machine Intelligence, 1(5):206–215, 2019.

[49] N. A. Rizvi et al. Cancer immunology. Mutational landscape determines sensitivity to PD-1 blockade in non-small cell lung cancer. Science(New York, N.Y.), 348(6230):124–128, April 2015.

[50] D. Dejardin et al. A composite decision rule of CD8+ t-cell density in tumor biopsies predicts efficacy in early-stage, immunotherapy trials. Clinical Cancer Research, 30(4):877–882, 2024.

[51] J. Liu et al. Cell cycle on the crossroad of tumorigenesis and cancer therapy. Trends in Cell Biology, 32(1):30–44, 2022.

[52] J. Wagner et al. Overexpression of the novel senescence marker β-galactosidase (glb1) in prostate cancer predicts reduced psa recurrence. PloS one, 10(4):e0124366, 2015.

[53] M. Ilic et al. Overexpression of Senescence-Associated Beta-Galactosidase (SA-B-GAL) asa Prognostic Marker of Invasive Breast Carcinoma. EABR. Experimental and Applied Biomedical Research, 26(1):39–52, January 2025.

[54] C. L. Cotarelo et al. Detection of Cellular Senescence Reveals the Existence of Senescent Tumor Cells within Invasive Breast Carcinomas and Related Metastases. Cancers, 15(6):1860, March 2023.

[55] M. L. B. Jr et al. Persistence of senescent prostate cancer cells following prolonged neoadjuvant androgen deprivation therapy. PLOSONE, 12(2):e0172048, February 2017. Publisher:Public Library of Science.

[56] A. M. Haugstetter et al. Cellular senescence predicts treatment outcome in metastasised colorectal cancer. British Journal of Cancer,103(4):505–509, August 2010. Publisher: NaturePublishing Group.

[57] K. He et al. Deep residual learning for image recognition. In Proceedings of the IEEE conference on computer vision and pattern recognition,pp. 770–778, 2016.

[58] A. Kondepudi et al. Foundation models for fast, label-free detection of glioma infiltration. Nature, pp. 1–7, 2024.

[59] R. Balestriero and Y. LeCun. Learning by reconstruction produces uninformative features for perception. arXiv preprint arXiv:2402.11337,2024.

[60] L. Farndale et al. Trident: Triple deep network training for privileged knowledge distillation in histopathology. arXiv preprint arXiv:2312.02111, 2023.

[61] L. Farndale et al. Synthetic privileged information enhances medical image representation learning. arXiv preprint arXiv:2403.05220, 2024.

[62] L. Farndale et al. Divide and conquer self-supervised learning for high-content imaging. arXiv preprint arXiv:2503.07444, 2025.

[63] N. Otsu. A threshold selection method from gray-level histograms. IEEE Transactions on Systems, Man, and Cybernetics, 9(1):62–66, 1979.

[64] S. Van der Walt et al. scikit-image: image processing in python. PeerJ, 2:e453, 2014.

[65] V. A. Traag et al. From louvain to leiden: guaranteeing well-connected communities. Scientific reports, 9(1):1–12, 2019.

[66] L. McInnes et al. UMAP: Uniform manifold approximation and projection for dimension reduction. arXiv preprint arXiv:1802.03426, 2018.

[67] O. Ronneberger et al. U-Net: Convolutional networks for biomedical image segmentation. In Medical Image Computing and Computer-Assisted Intervention – MICCAI 2015, volume 9351 of Lecture Notes in Computer Science, pp. 234–241. Springer, 2015.

[68] W. E. Johnson et al. Adjusting batch effects in microarray expression data using empirical Bayes methods. Biostatistics, 8(1):118–127, 2007.

[69] F. A. Wolf et al. SCANPY: large-scale single-cell gene expression data analysis. Genome Biology, 19(1):15, 2018.

[70] F. Pedregosa et al. Scikit-learn: Machine Learning in Python. Journal of Machine Learning Research, 12:2825–2830, 2011.

[71] D. D. Lee and H. S. Seung. Learning the parts of objects by non-negative matrix factorization. Nature, 401(6755):788–791, 1999.

[72] E. B. Garon et al. Pembrolizumab for the treatment of non-small-cell lung cancer. New England Journal of Medicine, 372(21):2018–2028, 2015.

[73] G. Palla et al. Squidpy: a scalable framework for spatial omics analysis. Nature methods, 19(2):171–178, 2022.

[74] M. Varrone et al. Cellcharter reveals spatial cell niches associated with tissue remodeling and cell plasticity. Nature genetics, 56(1):74–84, 2024.

[75] C. Davidson-Pilon. lifelines: survival analysis in Python. Journal of Open Source Software, 4(40):1317, 2019.

[76] F. E. Harrell et al. Evaluating the yield of medical tests. JAMA, 247(18):2543–2546, 1982.

[77] M. B. Amin et al., editors. AJCC Cancer Staging Manual. Springer, New York, 8th edition, 2017.

[78] A. L. Moreira et al. A grading system for invasive pulmonary adenocarcinoma: a proposal from the International Association for the Study of Lung Cancer pathology committee. Journal of Thoracic Oncology, 15(10):1599–1610, 2020.

[79] A. Lugli et al. Recommendations for reporting tumor budding in colorectal cancer based on the International Tumor Budding Consensus Conference (ITBCC) 2016. Modern Pathology, 30(9):1299–1311, 2017.

[80] K. Klintrup et al. Inflammation and prognosis in colorectal cancer. European Journal of Cancer, 41(17):2645–2654, 2005.

[81] J. H. Park et al. The relationship between tumour stroma percentage, the tumour microenvironment and survival in patients with primary operable colorectal cancer. Annals of Oncology, 25(3):644–651, 2014.

[82] F. E. Harrell et al. Multivariable prognostic models: issues in developing models, evaluating assumptions and adequacy, and measuring and reducing errors. Statistics in Medicine, 15(4):361–387, 1996.

[83] M. Ester et al. A density-based algorithm for discovering clusters in large spatial databases with noise. In Proceedings of the Second International Conference on Knowledge Discovery and DataMining (KDD-96), pp. 226–231, 1996.

[84] H. Edelsbrunner et al. On the shape of a set of points in the plane. IEEE Transactions on Information Theory, 29(4):551–559, 1983.

[85] J. A. Bull et al. Extended correlation functions for spatial analysis of multiplex imaging data. Biological Imaging, 4:e2, 2024.

[86] B. Delaunay. Surla sphére vide. Bulletin de l’Académie des Sciences de l’URSS, Classe des sciences mathématiques et naturelles, 6:793–800, 1934.

[87] C. Stringer et al. Cellpose: a generalist algorithm for cellular segmentation. Nature Methods, 18(1):100–106, 2021.

[88] E. C. Fieller et al. Tests for rank correlation coefficients. I. Biometrika, 44(3/4):470–481, 1957.

[89] Y. Benjamini and Y. Hochberg. Controlling the false discovery rate: a practical and powerful approach to multiple testing. Journal of the Royal Statistical Society: Series B (Methodological), 57(1):289–300, 1995.

[90] H. L. Williams et al. Spatial resolution of transcriptomic plasticity states underpinning lethal morphologies in lung adenocarcinoma, June 2024.Pages: 2024.06.11.598228 Section: New Results.

[91] M. I. Love et al. Moderated estimation of fold change and dispersion for RNA-seq data with DESeq2. Genome Biology, 15(12):550, December 2014.

[92] G. Korotkevich et al. Fast gene set enrichment analysis, February 2021. Pages: 060012 Section: New Results.

[93] S. Hänzelmann et al. GSVA: gene set variation analysis for microarray and RNA-Seq data. BMC Bioinformatics, 14(1):7, January 2013.

[94] M. J. Goldman et al. Visualizing and interpreting cancer genomics data via the Xena platform.Nature Biotechnology, 38(6):675–678, June 2020. Publisher: Nature Publishing Group.

