## Supplementary Figures for "Self-Supervised AI Reveals a Hidden Landscape of Prognostic Spatial Patterns in Multiplex Immunofluorescence Images"

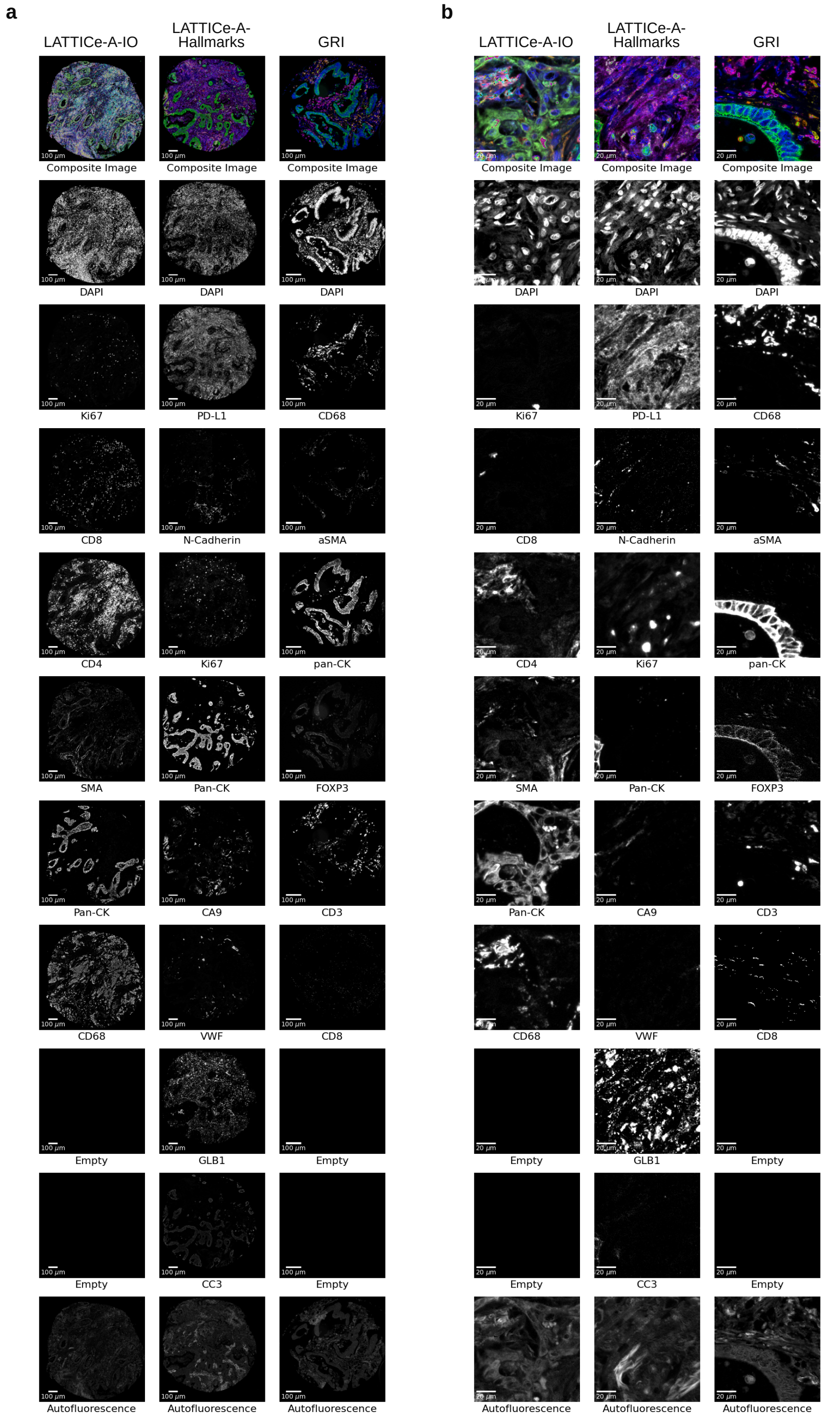


Supplementary Figure S1: **Example cores and tiles from the three multiplex panels.** Whole cores (a) and tiles (b) from LATTICe-A-IO, LATTICe-A-Hallmarks and GRI: composite pseudocolour image (top) and each channel in greyscale, including autofluorescence; "Empty", unused channel. Each marker highlights a distinct cell type or tissue compartment.

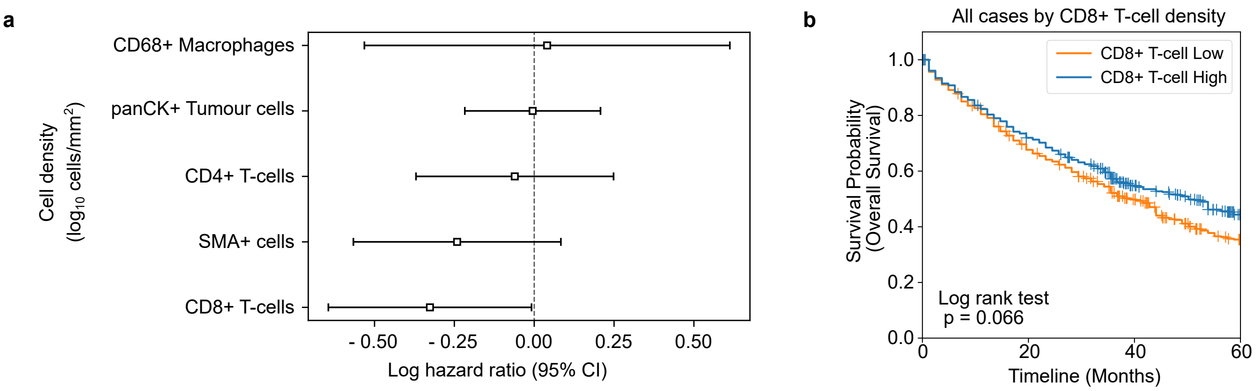


Supplementary Figure S2: **Cell-type densities alone carry little prognostic information in LATTICe-A-IO.** (a) Forest plots of multivariate Cox proportional hazards models of overall survival by $\log_{10}$ cell densities of basic phenotypes from the LATTICe-A-IO panel. (b) Kaplan Meier curve of the whole cohort dichotomised by the median CD8$^+$ T-cell density of the population. CD8$^+$ T-cell high n=461; CD8$^+$ T-cell low, n=461

| Channel | Lower threshold | Higher threshold |
|---|---|---|
| CD68 | 2.7 | 41.6 |
| $\alpha$SMA | 0.24 | 18.15 |
| panCK | 0.44 | 20.9 |
| FOXP3 | 0.3 | 5.31 |
| CD3 | 1.6 | 30.58 |
| CD8 | 0.3 | 13.51 |

Table S1: Threshold bounds (fluorescence intensity) for cell phenotyping in the GRI panel.

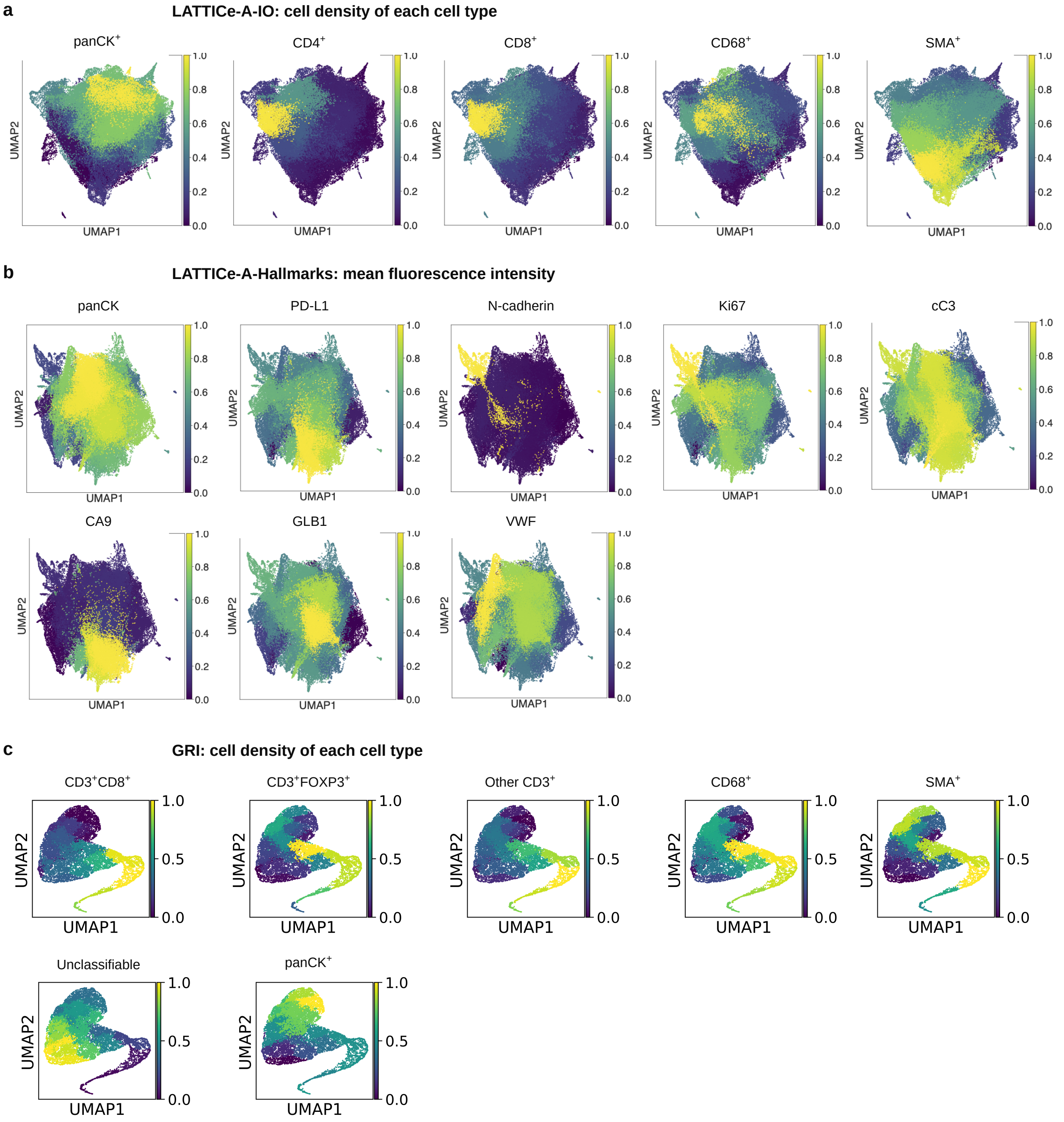


Supplementary Figure S3: **The SSL embeddings are organised by cell composition and marker expression.** UMAPs of the tile embeddings of LATTICe-IO (a), LATTICe-A-Hallmarks (b) and GRI (c), as in Figures 3a, b and 5a, coloured by the scaled mean density of each cell type (IO, GRI) or the scaled mean intensity of each marker (Hallmarks). Each cell type or marker occupies a coherent region of the embedding.

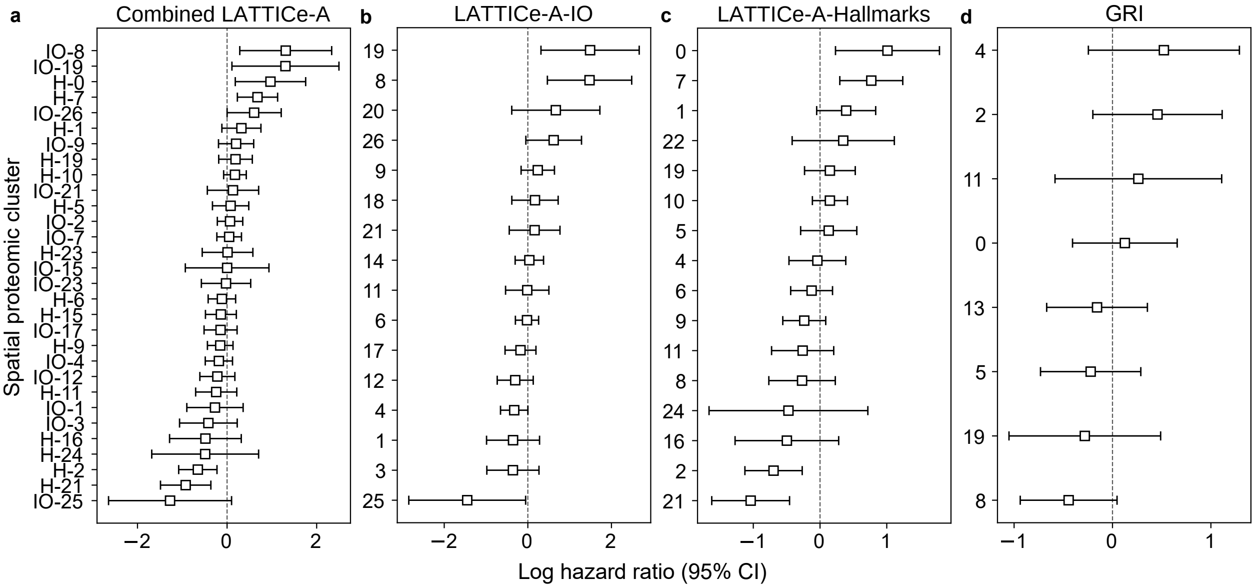


Supplementary Figure S4: **Hazard ratios of individual SPCs in each panel.** Forest plots showing cluster associations with survival for each panel. In the cluster labels, “IO” and “H” indicate the LATTICe-A-IO and LATTICe-A-Hallmarks panels respectively. Error bars are 95% confidence intervals.

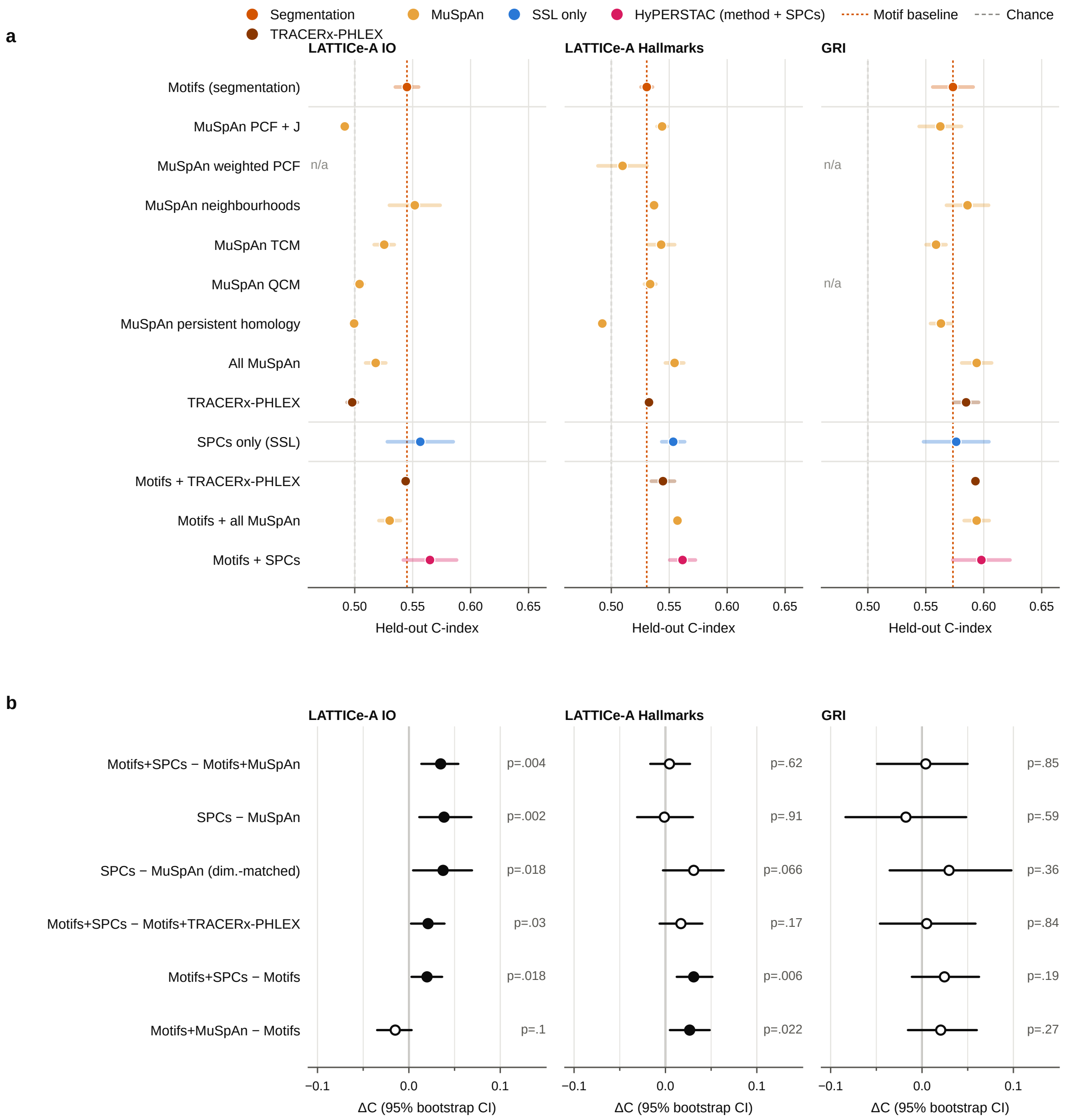


Supplementary Figure S5: **Full benchmark: SPCs add to established methods.** (a) Held-out C-index of every feature set (dot, mean of the two folds; bar, range; dotted orange line, motif baseline; dashed line, chance), coloured by family: segmentation motifs, TRACERx-PHLEX, MuSpAn families, SPCs alone (SSL only) and HyPERSTAC (method + SPCs). n/a, not computable for that panel. (b) Paired differences in held-out C-index (ΔC; 95% bootstrap CI; filled, CI excludes 0; right, $p$). "Dim.-matched", MuSpAn reduced by PCA to the number of SPCs.

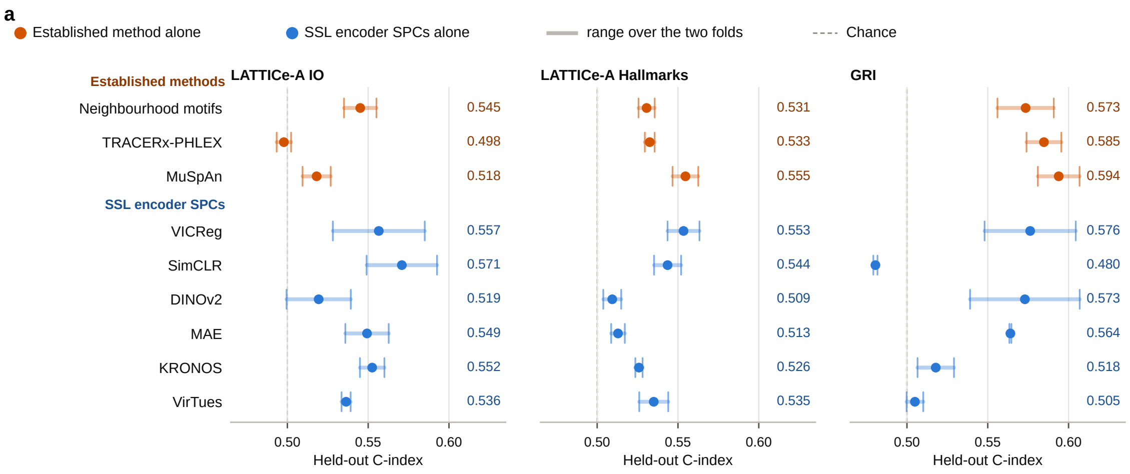


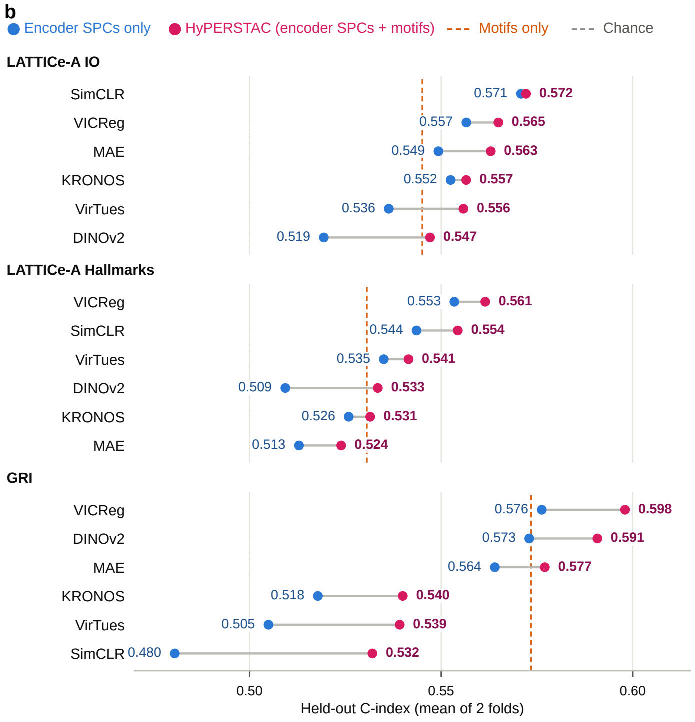


**c**

| Encoder | IO | Hallmarks | GRI | Avg. rank |
|---|---|---|---|---|
| **VICReg** | **2 (0.565)** | **1 (0.561)** | **1 (0.598)** | **1.3** |
| SimCLR | 1 (0.572) | 2 (0.554) | 6 (0.532) | 3.0 |
| DINOv2 | 6 (0.547) | 4 (0.533) | 2 (0.591) | 4.0 |
| MAE | 3 (0.563) | 6 (0.524) | 3 (0.577) | 4.0 |
| VirTues | 5 (0.556) | 3 (0.541) | 5 (0.539) | 4.3 |
| KRONOS | 4 (0.557) | 5 (0.531) | 4 (0.540) | 4.3 |

Rank (C) of encoder SPCs + motifs; 1 = highest.

Supplementary Figure S6: **Adding SPCs to motifs helps most with encoders trained on the study images.** (a) Held-out C-index of each feature set alone (dot, mean of the two folds; bar, range): established methods (orange) and SPCs from each encoder (blue). (b) Held-out C-index of SPCs alone (blue) and HyPERSTAC (SPCs + motifs; magenta) for all six encoders; dashed orange line, motifs alone. SPCs improved C-index over motifs alone for all models. (c) Encoder ranks for HyPERSTAC, averaged over the three cohorts: VICReg ranks first.

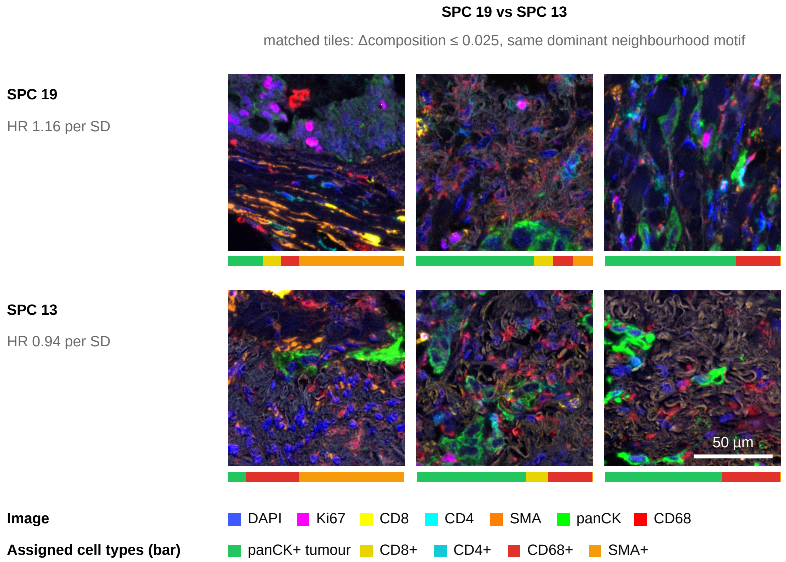


Supplementary Figure S7: **Pre-specified composition-matched tile pair (SPC 19 vs SPC 13).** As in Figure 2d, for SPC 19 (training-fold HR 1.16 per s.d.) and SPC 13 (HR 0.94), the pair chosen before any outcome comparison (mean compositions within total-variation distance 0.10, opposite training-fold HRs, largest difference in log HR). The three tile pairs, all six tiles from different patients, have composition distance $\leq 0.025$ and the same dominant motif. Scale bar, 50 $\mu$m.

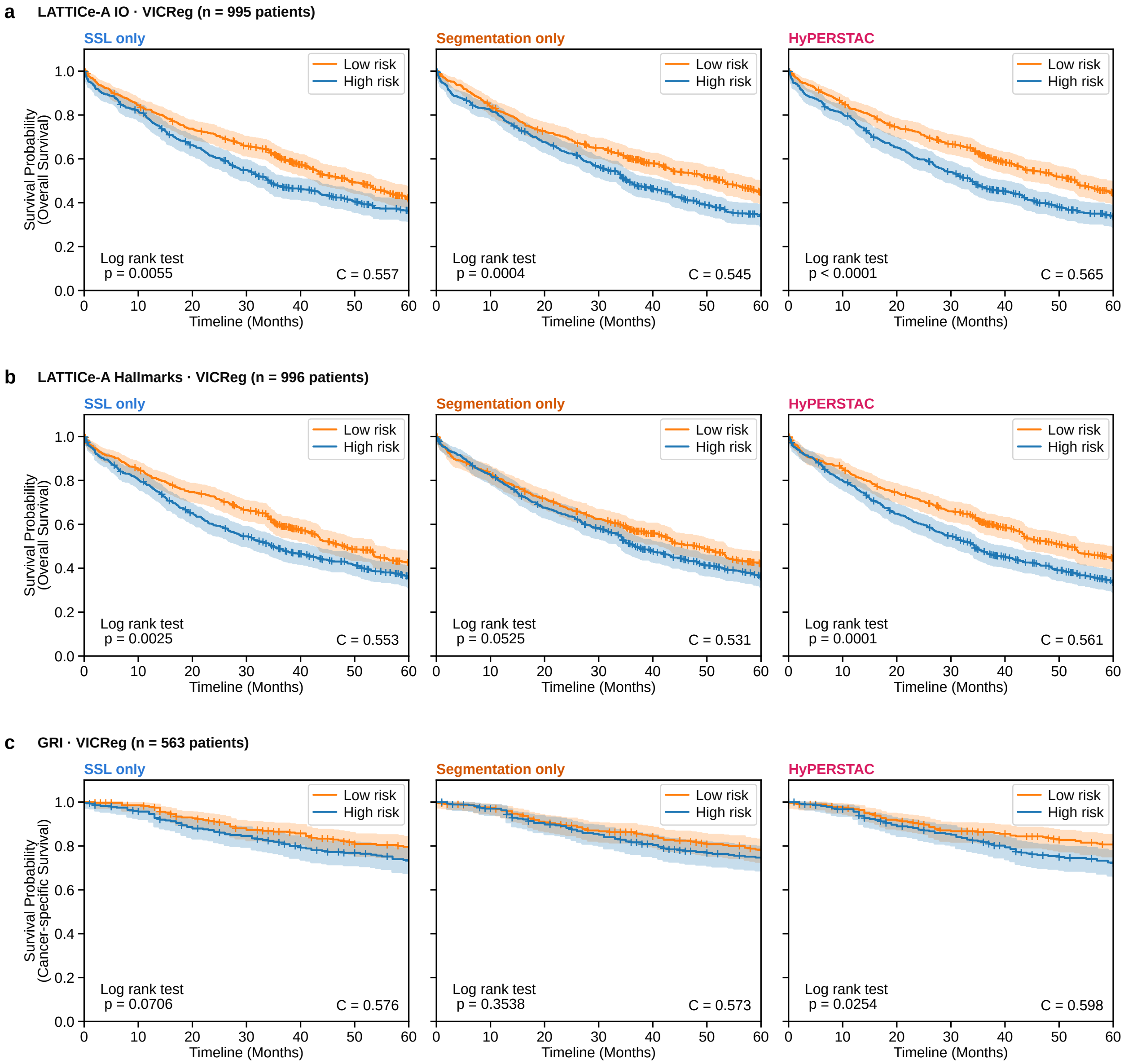


Supplementary Figure S8: **HyPERSTAC stratifies held-out patients in all three cohorts.** Kaplan–Meier curves for SSL only (VICReg SPCs), segmentation only (neighbourhood motifs) and HyPERSTAC (motifs + SPCs), with held-out patients of both folds pooled and risk cut-points set on training patients. (a) LATTICe-A-IO ($n = 995$; overall survival). (b) LATTICe-A-Hallmarks ($n = 996$; overall survival). (c) GRI ($n = 563$; cancer-specific survival). Log-rank p, lower left; C-index, lower right; shading, 95% CI; truncated at 60 months.

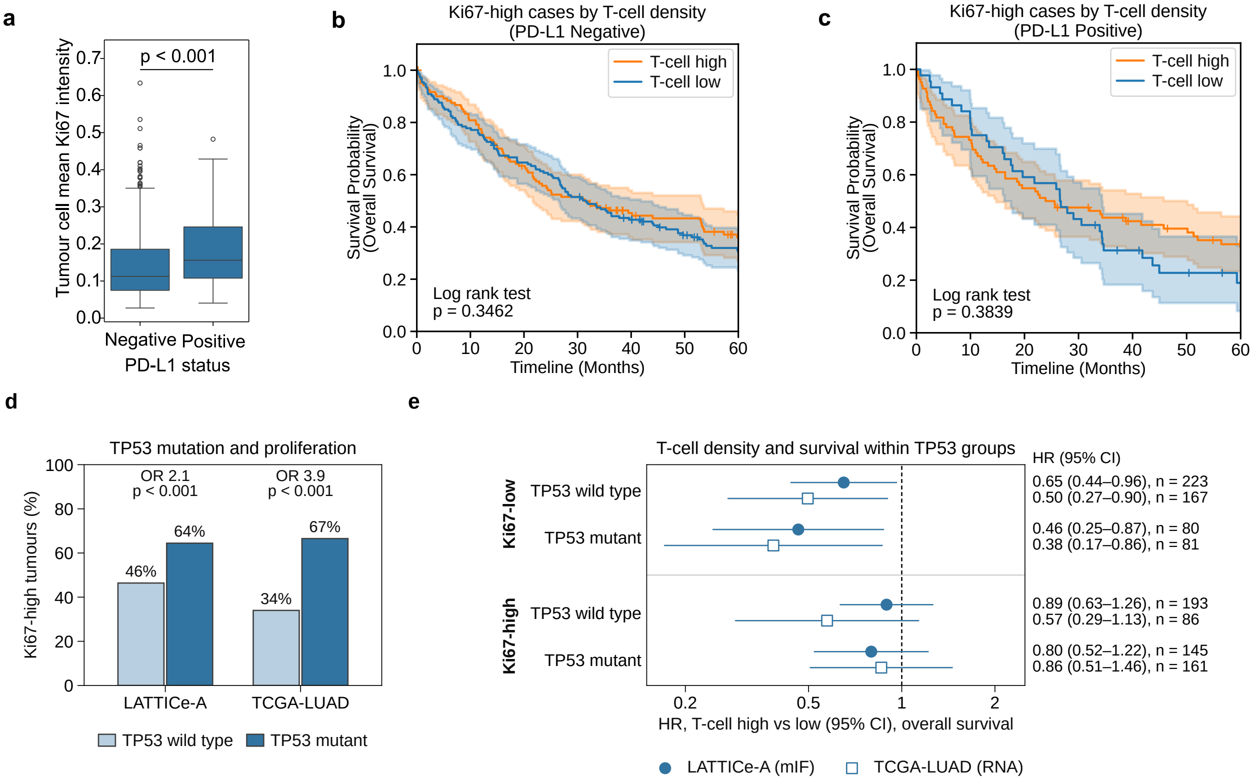


Supplementary Figure S9: **The lack of T-cell benefit in Ki67-high tumours does not depend on PD-L1 status.** (a) Tumour-cell Ki67 intensity is higher in PD-L1$^+$ than PD-L1$^-$ tumours (Mann–Whitney $p < 0.001$). (b, c) Overall survival of Ki67-high tumours by T-cell density in PD-L1$^-$ (b; $n = 275$; log-rank $p = 0.35$) and PD-L1$^+$ (c; $n = 126$; $p = 0.38$) tumours: T-cell density is not associated with survival in either. Shading, 95% CI; +, censored. (d) Share of Ki67-high tumours by TP53 status in LATTICe-A ($n = 641$) and TCGA-LUAD ($n = 495$; Fisher test): TP53-mutant tumours are more often highly proliferative. (e) HR (95% CI) for T-cell high vs low within each Ki67 $\times$ TP53 group (filled circles, LATTICe-A; open squares, TCGA-LUAD): T cells are associated with longer survival in Ki67-low tumours whether TP53 is wild type or mutant, and not in Ki67-high tumours of either type, so the pattern follows proliferation rather than TP53 status. Overall survival truncated at 60 months; associative.

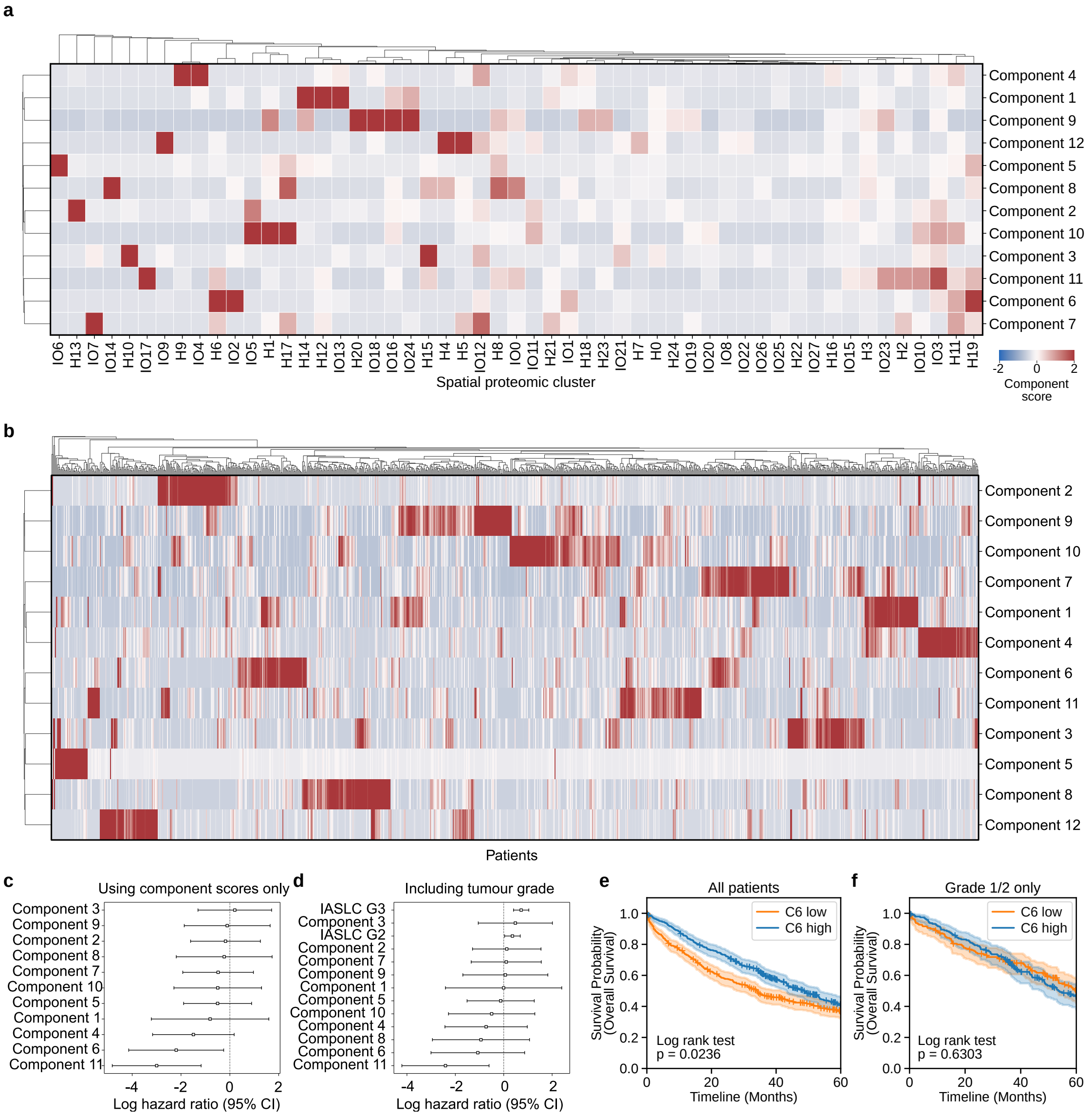


Supplementary Figure S10: **Cross-panel NMF components and their survival associations.** (a) Definitions of the 12 components: heatmap of the NMF component scores of each spatial proteomic cluster (IO SPCs prefixed IO, Hallmarks SPCs prefixed H). A high score indicates a strong contribution of that SPC to that component's score. Rows (components) and columns (SPCs) are hierarchically clustered. Colour bar, component score, $-2$ to 2. (b) Heatmap of LATTICe-A patients and component scores, as z-scores. (c) Forest plot showing the association between component scores and overall survival from a multivariate Cox proportional hazards model. (d) Forest plot showing the association between component scores and overall survival from a multivariate Cox proportional hazards model, when also including IASLC grade as a feature. (e, f) Overall survival by component 6 (pre-specified median split) in all patients (e; $n = 906$; log-rank $p = 0.024$) and grade 1–2 tumours (f; $n = 384$; $p = 0.63$): the association is absent in low-grade disease (grade 3 in Figure 6g). Cohort as Figure 6g; associative analyses without cross-validation. Shading, 95% CI; +, censored.

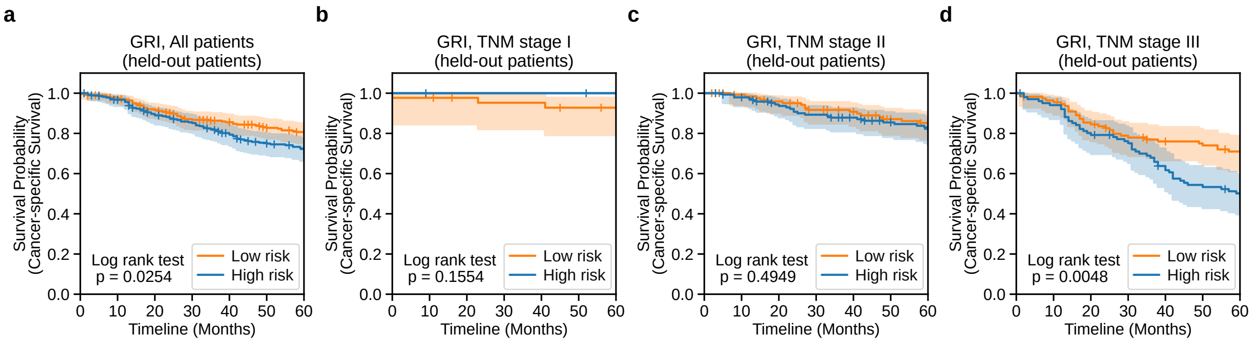


Supplementary Figure S11: **The HyPERSTAC risk score in GRI by TNM stage: separation is concentrated in stage III.** Cancer-specific survival of held-out GRI patients split into high (blue) and low (orange) risk by the HyPERSTAC score of Figure 5b, with risk groups defined once in the full cohort and then restricted to each stage (no within-stage cut-point). (a) All patients ($n = 563$; log-rank $p = 0.025$). (b) Stage I ($n = 71$; only 3 deaths; $p = 0.16$). (c) Stage II ($n = 276$; $p = 0.49$). (d) Stage III ($n = 215$; $p = 0.0048$; as Figure 5b). Shading, 95% CI; +, censored; truncated at 60 months.

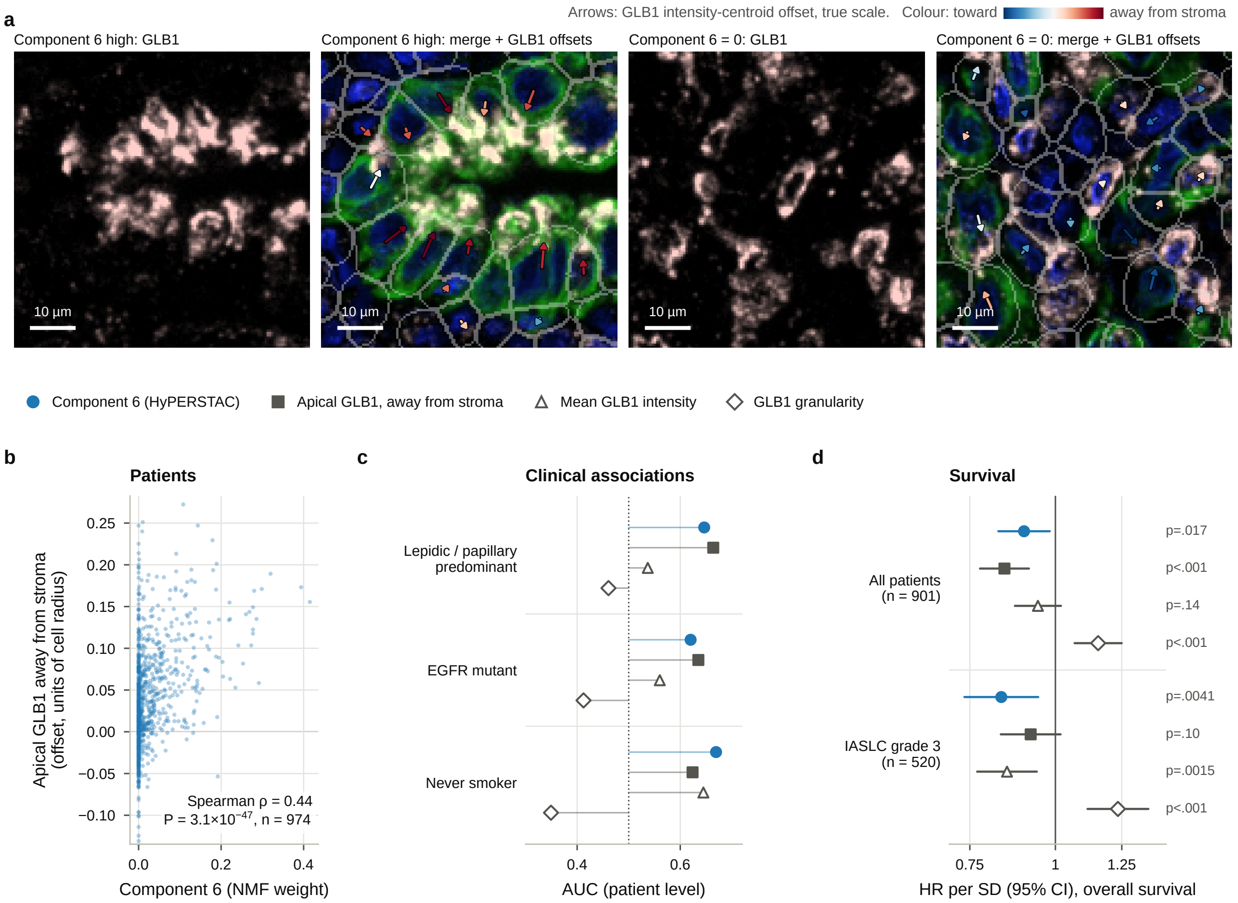


Supplementary Figure S12: **HyPERSTAC component 6 (apical GLB1) can be defined post-hoc through segmentation-based features.** (a) Per-cell GLB1 offsets (arrows, true scale; red, away from stroma; blue, towards) in a component 6-high core (left pair; as Figure 6i) and a component 6 = 0 core with GLB1 at least as bright (right pair); each pair shows GLB1 alone and a merge (colours as Figure 6b: DAPI blue, panCK green, GLB1 pink; grey, cell outlines). Offsets are of similar size in both cores (median 1.9 vs 1.8 $\mu$m), but only in the component 6-high core do they point away from stroma: the cores differ in the direction of GLB1, not its displacement. 65 $\mu$m windows; scale bars, 10 $\mu$m. (b) Per-patient apical GLB1 offset against component 6 ($n = 974$; Spearman $\rho = 0.44$). (c) Discrimination (AUC) of lepidic/papillary growth, *EGFR* mutation and never-smoking by component 6 (blue) and the explicit GLB1 features (grey; square, apical offset as in Figure 6j); AUC $< 0.5$, feature lower in the positive group. (d) HR per s.d. for overall survival in all patients ($n = 901$) and IASLC grade 3 ($n = 520$): the apical offset matches component 6 overall but is weaker in grade 3. The explicit features were designed after component 6 was identified.

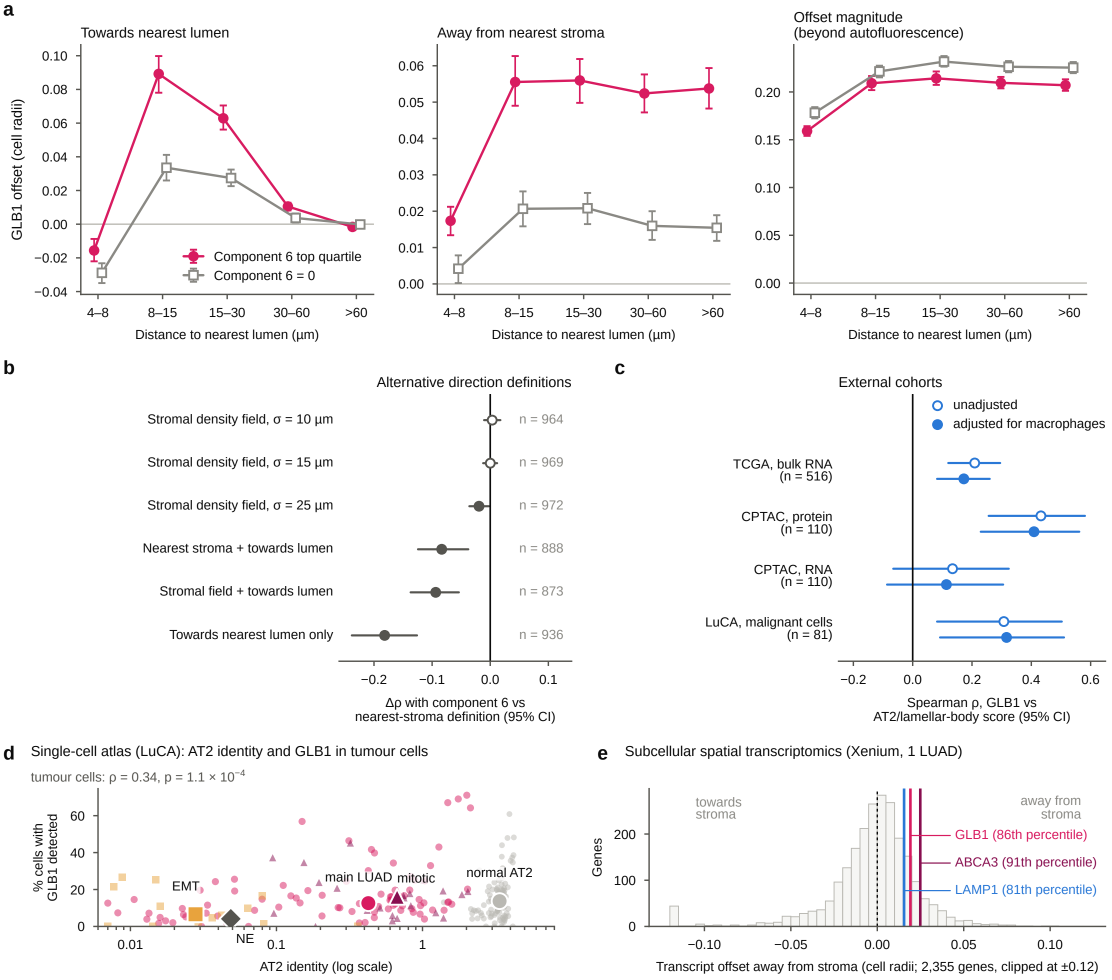


Supplementary Figure S13: **GLB1 polarity differs in direction rather than amount, and GLB1 tracks the alveolar type II (AT2) program and is polarised away from stroma in external data.** (a) GLB1 offset direction of GLB1$^+$ tumour cells (cell radii; autofluorescence-referenced) by distance to the nearest lumen, for component 6 top-quartile (magenta) and component $6 = 0$ patients (grey): towards the lumen, away from the nearest stromal cell, and magnitude beyond the autofluorescence offset. Mean of patient means with bootstrap 95% CI; exploratory. (b) Alternative polarity directions: difference in Spearman correlation with component 6 relative to the nearest-stromal-cell definition ($\rho = 0.44$, $n = 974$); paired bootstrap 95% CI; filled, CI excludes 0. (c) Spearman correlation between GLB1 and an AT2/lamellar-body score in TCGA-LUAD RNA, CPTAC LUAD protein and RNA, and LuCA malignant LUAD cells (patient pseudobulk), unadjusted (open) and adjusted for a macrophage score (filled; CPTAC also for purity); Fisher-$z$ 95% CIs. (d) LuCA: AT2 identity (log scale) against the percentage of cells with *GLB1* detected, per patient and cell state ($\geq$20 cells; small markers) and per-state medians (large markers); normal AT2 cells for reference; two groups with AT2 identity $\leq$0.007 drawn at the axis edge. (e) Xenium Prime 5K lung adenocarcinoma (one patient): mean offset of each gene's transcripts from the cell's transcript centroid, away from the nearest stromal cell, over epithelial cells within 15 $\mu$m of stroma (2,355 genes; clipped at $\pm$0.12); lines mark *GLB1*, *ABCA3* and *LAMP1* with percentiles. The most positive genes are apical mucins and the most negative stromal matrix genes, probably from neighbouring stromal cells. Methods give gene sets and data sources.

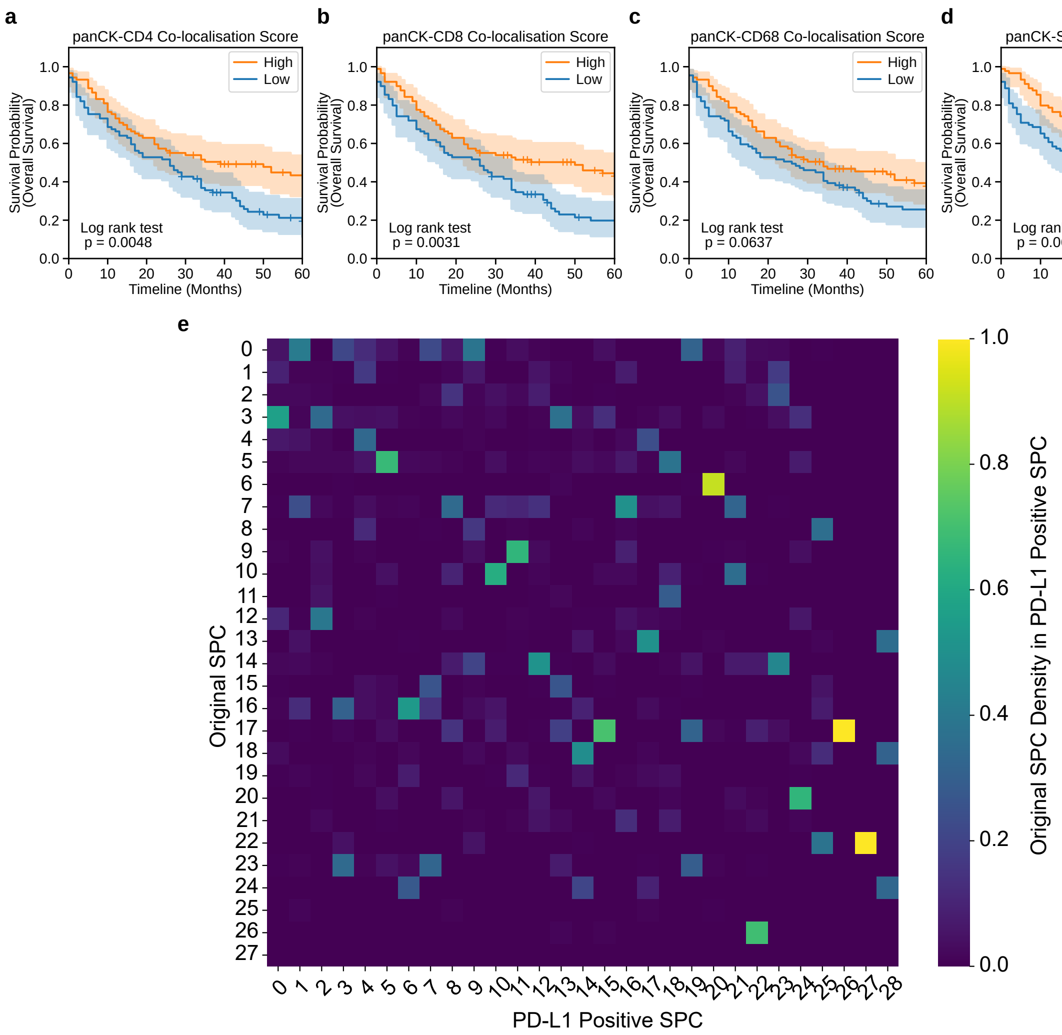


Supplementary Figure S14: **PD-L1$^+$ patients stratified by spatial co-localisation score of tumour cells and CD4$^+$ T-cells.** (a–d) Overall survival of PD-L1$^+$ patients split at the median co-localisation score of tumour cells with CD4$^+$ T cells (a; log-rank $p = 0.0048$), CD8$^+$ T cells (b; $p = 0.0031$), CD68$^+$ macrophages (c; $p = 0.064$) and SMA$^+$ cells (d; $p = 0.060$); $n = 89$ per group; associative. More tumour–T-cell contact goes with longer survival. Shading, 95% CI; +, censored. (e) Fraction of each PD-L1$^+$SPC's tiles (columns) that belong to each original SPC (rows): most PD-L1$^+$SPCs map onto one or a few original SPCs; for example, PD-L1$^+$SPC 26 maps onto SPC 17.

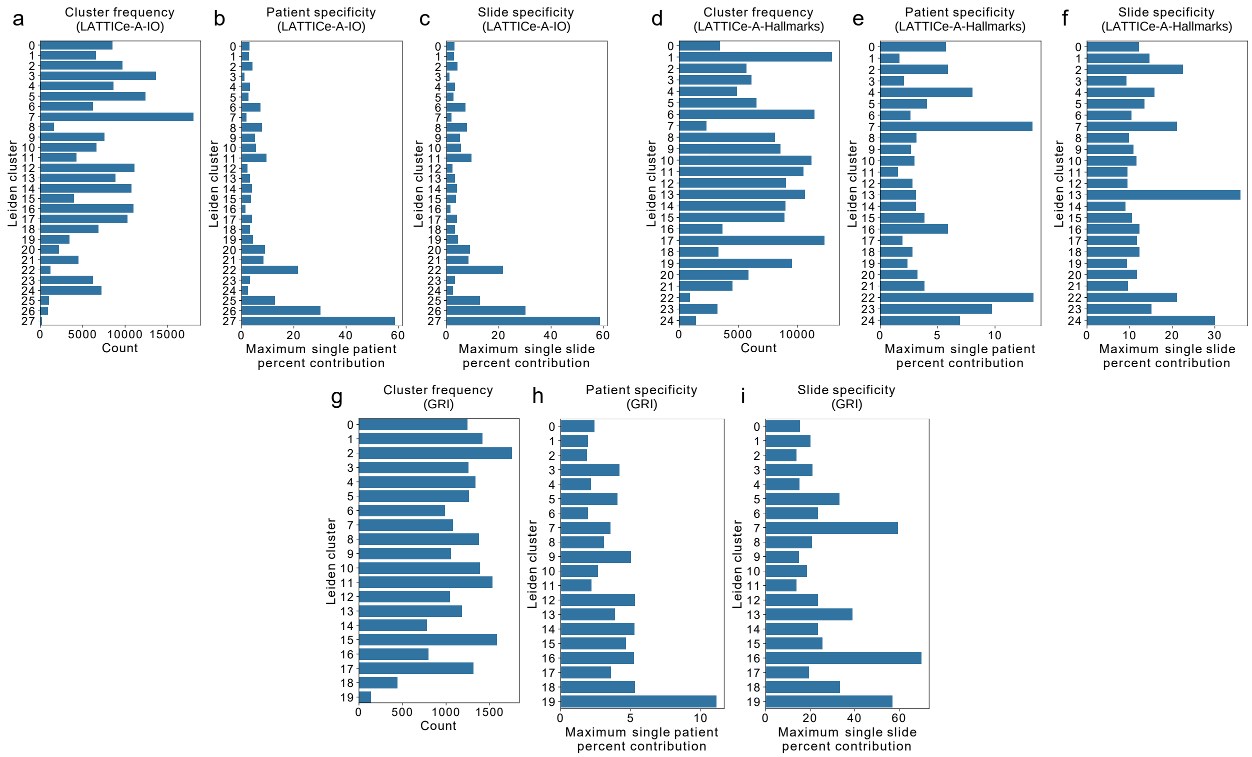


Supplementary Figure S15: **SPCs recur across many patients and slides rather than reflecting a single patient or batch.** (a, d, g) Raw cluster frequency in LATTICe-A-IO (a), LATTICe-A-Hallmarks (d) and GRI (g) cohorts. (b, e, h) Patient specificity measures for the clusters discovered from LATTICe-A-IO (b), LATTICe-A-Hallmarks (e) and GRI (h) calculated as the largest fraction of instances of a cluster from a single patient. (c, f, i) Slide specificity measures for the clusters discovered from LATTICe-A-IO (c), LATTICe-A-Hallmarks (f) and GRI (i) calculated as the largest fraction of instances of a cluster from a single slide.
